# The effectiveness of two new *Metarhizium brunneum* formulations against wireworms in potato cultivation

**DOI:** 10.1101/2024.12.27.630518

**Authors:** Maximilian Paluch, Tanja Seib, Dietrich Stephan, Eckhard Immenroth, Jörn Lehmhus

## Abstract

In the absence of officially approved agrochemicals for the use against wireworms (*Agriotes* spp.) in German potato cultivation, the entomopathogenic fungus genus *Metarhizium* has been regarded as promising biocontrol agent. However, previous mycoinsecticide formulations of *M. brunneum* could not achieve consistent effectiveness in field use. Within the scope of the project AgriMet, the effectiveness of two new formulations of *M. brunneum* was evaluated over three years in the field, greenhouse and laboratory. The application of a soil granule and/or a wettable powder (dry product) of *M. brunneum* during potato planting aimed to reduce potato tuber damage by controlling wireworms. During field trials in the Lower Saxony (Germany), the combination of both formulations led to an effectiveness between 1 % and 13.6 % compared to the untreated control. The single application of the AgriMet-Granule and the AgriMet-Dry Product was less successful. The standardised greenhouse set-up in pots with *A. obscurus* indicated that an application rate of at least 300 kg ha^-1^ of the AgriMet-Granule, respectively 1×10^13^ conidia ha^-1^ of the AgriMet-Dry Product was necessary to effectively reduce tuber damage (44-54 %), while surprisingly no larval mortality was recorded. The laboratory experiments showed, that only the AgriMet-Dry Product provided a lethal potential against wireworms, whereas the AgriMet-Granule showed deficits in terms of product quality. However, the effectiveness of the AgriMet-Dry Product against *A. obscurus*, *A. sputator* and *A. lineatus* resulted in significant differences in mortality indicating a species-specific virulence of the *M. brunneum* isolate used. Additional influencing factors became apparent through analysing multiple soil parameters at the respective field site such as temperature, moisture, colony-forming units of *Metarhizium* spp. and wireworm density, underlining the complexity of wireworm control using a microbial control agent. Especially the low soil temperatures during spring application pose a serious challenge to the development of an effective control strategy against wireworms.

## Introduction

Wireworms are the larvae of click beetles (Coleoptera: Elateridae) and a major pest of a variety of crops including sugar beet, cereals and potatoes. Potatoes (*Solanum tuberosum*) are particularly affected by wireworms, as the soil-dwelling larvae reduce quality rather than yield and tuber damage is usually not detected until harvest (Parker & Howard 2001). Tuber damage consists of narrow tunnels in the tuber flesh as wireworms feed their way into potato tubers in search of food and moisture (Miles 1942, Parker & Howard 2001, Vernon & Van Herk 2013b, Traugott et al. 2015). The typical small holes on the surface provide entry points for the fungal plant pathogen *Rhizoctonia solani* and resulting “drycore” symptoms at the tubers cause serious economic losses (Keiser et al. 2012). Potato lots with significant wireworm damage are not marketable as fresh potatoes because consumers demand high quality without any blemish (Willersinn et al. 2015). Vernon & Van Herk (2013a) suggested that a threshold of less than 5 % of total yield damaged through wireworms would generally meet most industry standards, making it challenging to grow potatoes for direct food consumption. In Germany, especially the species *Agriotes obscurus*, *A. sputator* and *A. lineatus* are a problem to potato cultivation (Lehmhus 2019). Larvae of the genus *Agriotes* spend 3-5 years in soil before pupating (Schepl & Paffrath 2010) posing a harm to the entire crop rotation (Myers et al. 2008). Due to the threat of wireworms to agronomic productivity, the control of these larval pests is of economic importance (Parker & Howard 2001).

Efficient control of wireworms has historically been achieved by the use of insecticidal chemicals such as Fipronil or Thiametoxam (Ritter & Richter 2013). However, the European Commission restricted the approval of these agrochemicals in most indications due to harmful effects on bees assessed by the European Food Safety Authority (EFSA 2013a, EFSA 2013b). These regulations left German potato growers without fast-acting tools against the soil-dwelling pests (BVL 2021). Limited wireworm control options enhance the research of sustainable and environmentally friendly alternatives to profit from the socioeconomic benefits of reducing wireworms in potato cultivation (Benjamin et al. 2018). Different authors reviewed various approaches to reduce wireworms within the integrated pest management (IPM), but stated fluctuating control success (Parker & Howards 2001; Barsics et al. 2013). The authors postulated that entomopathogenic fungi are most promising for developing efficient control strategies against wireworms and efforts to use the species *Metarhizium brunneum* have been intensively pursued (Ericsson et al. 2007, Kabaluk & Ericsson 2007, Reddy et al. 2014, Brandl et al. 2017). The entomopathogenic fungal genus *Metarhizium* is well-studied and active against several agricultural pests such as locust or termites. The mode of action is based on the adhesion of the infectious units, the conidia, to the host cuticle initiating the lethal infection cycle (Zimmermann 2007). In vitro studies confirmed the lethal potential of *Metarhizium* spp. against wireworms of the genus *Agriotes*. The most efficient strains caused up to 73-100 % mortality approximately 8 weeks after exposure, depending on the wireworm species tested (Ansari et al. 2009, Eckard et al. 2014). In light of the pathogenicity of fungal isolates against the larval pests, formulation and commercialisation as biopesticides, more precisely mycoinsecticides, for the use in potato cultivation seems conceivable. However, the challenging task developing an effective mycoinsecticides is to mass-produce stable fungal propagules and to formulate them in most appropriate manner for soil application (Zaki et al. 2020).

The formulation of the biocontrol agent *M. brunneum* as mycoinsecticide is one of the most important factors for its commercial success, as it preserves the organism, deliver it to the target site while enhancing its biological activity (Jones & Burges 1998). Two types of formulations are suitable for the practical implementation: suspensions and dry products (Rhodes 1993). Since the moisture content of fungal conidia affects their storage characteristics and thus important product requirements (Moore et al. 1996), formulation of entomopathogenic fungi as dry products is preferred. Dry products include wettable powders and granules (Jones & Burges 1998). Latter contain a synthetic or organic carrier in which the fungus is incorporated or attached on the outer surface. The carrier can serve as nutrient source for proliferation and increase its persistence after application, while the distribution at the target site is determined by the size of the carrier in combination with the application rate (Goss et al. 1996). Wettable powders consist of powdered fungal propagules miscible in water, which acts as a carrier. The spray mist containing the fungal propagules is distributed evenly, able to hit target pests directly by droplets or colonise organic material in the application area (Silva et al. 2015). Both granules and wettable powders are suitable mycoinsecticide formulations for the application during potato planting, as the technical requirements at common potato-planting machines are given. Since the granule spreader and the spray device used for this purpose function independently of each other, individual and combined use of a granule and wettable powder is conceivable (GRIMME Landmaschinenfabrik GmBH & Co. KG, Damme, Germany).

Within the project AgriMet, a granule and a wettable powder based on the entomopathogenic fungus *M. brunneum* were developed for the application during potato planting. Both formulations are supposed to reduce tuber damage by the control of wireworms. Although the pathogenicity of the used isolate has been tested in the laboratory, no results about the effectiveness of the AgriMet formulations against wireworms exists so far. The AgriMet-Granule consists of an autoclaved millet grain as carrier, which provides the required hardness and grain size for production and application (Stephan et al. 2020). Furthermore, autoclaved millet attracted wireworms of the genus *Agriotes* in an olfactrometric bioassay and is therefore suitable as bait for the “attract-and-kill” technique. The AgriMet-Dry Product is a wettable powder of *M. brunneum* that is soluble in tap water for a spray application of a defined amount of conidia directly into the potato ridge. The biological activity of both the AgriMet-Granule and the AgriMet Dry product should be provided by the formation of infectious conidia after application in soil. Since the release of the entomopathogenic fungus aims at its proliferation and establishment for an extended period, but not permanently, the control strategy represents inoculative biocontrol (Eilenberg et al. 2001). The desired control strategy assumes that fungal reproduction is possible under the abiotic and biotic soil conditions in the potato ridge. However, although comparable granule mycoinsecticide formulations against wireworms have been tested in the field, there are no data relating soil temperature and moisture to the control success (Humbert et al. 2017, Brandl et al. 2017, Sharma et al. 2020). Thus, an important aspect for the interpretation of inoculative control strategies against wireworms is missing and may explain the fluctuating effectiveness in field use.

In this study, the effectiveness of two new formulations of *M. brunneum* against wireworms for the application in potato cultivation was evaluated in more detail. First, the reduction of tuber damage through the application of the AgriMet-Granule and AgriMet-Dry Product was investigated in field trials from 2018-2020. Standardised bioassays in the laboratory and greenhouse provided information on the required application rate and the host spectrum of the mycoinsecticide formulations.

Additionally, parameters like soil temperature and water content, wireworm species composition and distribution at each field site were recorded. To verify fungal proliferation of the inoculative control strategy, the *Metarhizium* spp. concentration in the field soil (colony-forming units) was determined throughout the vegetation period after application of the formulations. In the end, the detailed evaluation of the tested AgriMet formulations for wireworm control in potato cultivation revealed important factors influencing their effectiveness.

## Material and Methods

### Metarhizium brunneum formulations

#### AgriMet-Granule

The *Metarhizium brunneum* isolate JKI-BI-1450, isolated in 2016 from an infected *Agriotes lineatus* beetle in Germany, was produced by liquid fermentation. The biomass, including submerged spores, hyphae and metabolites, was coated on autoclaved millet of the species *Seteria italic* (Alnatura Produktions-und Handels GmbH, Darmstadt, Germany) using a fluidised bed dryer. 1 kg AgriMet-Granule contained 4.5 g of the dry fungal biomass. The AgriMet-Granule was dry and dust-free indicated by the Heubach test of Immenroth, which resulted in 0.1-0.2 mg of abrasion per 100 g AgriMet-Granule. Comparable thresholds of the Heubach test for seed dressings are 2 mg abrasion for wheat or 9 mg abrasion for sugar beet (personal communication, Immenroth, 2020, JKI Institute for Application Techniques in Plant Protection, Braunschweig, Germany). The fungus was still active due to the low thermal load during the production process. The AgriMet-Granule was provided by Tanja Bernhardt (JKI Institute for Biological Control, Darmstadt, Germany). To ensure the functionality, the granule was stored at 5 °C and in darkness for no longer than eight weeks.

#### AgriMet-Dry Product

The AgriMet-Dry Product was a powder soluble in tap water for a spray application. Mass-production of the *M. brunneum* isolate JKI-BI-1450 was carried out by liquid fermentation. The fungal biomass was dried by lyophilisation and ground with a hand mortar to receive a *M. brunneum* powder with a concentration of 1.25×10^9^ spores g^-1^ according to the manufacturer. The AgriMet-Dry Product was provided by Marianne Buschke (ABiTEP GmBH, Berlin, Germany).

#### ATTRACAP®

ATTRACAP® was commercially available under an emergency authorisation (2018, 2019, 2020) and used as reference product. The recommended application rate of 30 kg ha^-1^ contained 1.2×10^10^ conidia. For the production of ATTRACAP® (BIOCARE Biologische Schutzmittel GmbH, Einbeck, Germany), aerial conidia of the *M. brunneum* isolate CB15-III are encapsulated with *Saccharomyces cerevisiae* and maize starch in wet spherical calcium alginate beads. The beads are dried using fluidised-bed drying. Under moist conditions the beads swell in the potato row and the yeast convert the encapsulated starch to CO_2_ (Humbert et al. 2017).

#### Quality control of the Metarhizium brunneum formulations

With each experiment, a simultaneous quality control was carried out to ensure the functionality of the formulations. The AgriMet-Granule and ATTRACAP® was examined by placing twenty grains or capsules on each of five Petri dishes with 1.5 % water agar. After 14 days at 25 °C in the dark, the number of grains or capsules with fungal growth of *Metarhizium brunneum* on the surface were determined to calculate the rate of outgrowth in percent. In addition, the sporulation of the *M. brunneum* isolate JKI-BI-1450 coated on the AgriMet-Granule was tested in different field soils at 10°C, 15 °C, 20 °C and 25 °C. Therefore, twenty grains of the AgriMet-Granule were placed on each of three Petri dishes with sterile 1.5 % water agar (Control) or untreated soil from the experimental field sites Borg and Suettdorf. The field soil was the same as that used for the analysis of nutritive substances. It was previously sieved (1.6 mm mesh) and moistened to ensure a homogenous substrate and comparability to the control. After 14 days of incubation at the respective temperature, ten grains per Petri Dish were washed with 0.1 % Tween80® and conidia were removed by vortexing to determine the concentration of conidia grain^-1^ with a haemocytometer. This part of the quality control was repeated three times. The AgriMet-Dry Product was verified by spreading 100 µl of a dilution in tap water on each of five Petri dishes filled with 1.5 % malt peptone agar. The plates were incubated for 14 days at 25 °C in darkness and the number of colony-forming units (CFUs) was determined to calculate the CFUs g^-1^ AgriMet-Dry Product. The identification of *M. brunneum* for each formulation was based on morphological criteria (Zimmermann 2007).

### Assessment of wireworm potato damage

Wireworm damage during field and greenhouse experiments was assessed following the European and Mediterranean Plant Protection Organization (EPPO) standard PP 1/46 on the conduct and evaluation of wireworm field trials. At harvest, 100 (field) or rather 10 (greenhouse) potato tubers per plot or pot were randomly sampled and categorised by classes based on the number of feeding holes (class 1: 0 holes, class 2: 1-2 holes, class 3: 3-5 holes, class 3: >5 holes). Feeding holes were defined as > 5 mm tunnels into the tuber. Assessment of wireworm potato damage in the field trials was conducted by the Lower Saxony Chamber of Agriculture.

### Wireworm breeding for the greenhouse and laboratory experiments

For a species-specific distinction, wireworms used in the greenhouse and laboratory experiments were taken from a laboratory breeding. *Agriotes obscurus, A. sputator* and *A. lineatus* beetles were caught in Wohld (52°18’11.0”N 10°41’11.6”E, Germany) and determined to species level using the identification key of Lohse (1979). Following the protocol of Kölliker et al. (2009) with minor modifications, larvae emerged from the breeding established at the JKI Institute for Plant Protection in Field Crops and Grassland (Braunschweig, Germany) and were stored dark at 5 °C in plastic boxes (18.3 x 13.6 x 6.4 cm, Baumann Saatzuchtbedarf, Waldenburg, Germany) with moist paper towel. 14 days before the start of an experiment wireworms were transferred into soil and stored dark at 15 °C for acclimatisation and nutrition with *Triticum aestivum* seeds (Cultivar: Primus, Deutsche Saatgutveredelung AG, Lippstadt, Germany). The larval stages were determined by measurement of the head width and varied between 83-154 mm by *A. obscurus,* 68-120 mm by *A. sputator* and 74-163 mm by *A. lineatus*.

### Effectiveness of the *Metarhizium brunneum* formulations in the field

#### Field sites

Field trials were conducted near Uelzen, Lower Saxony, the region with the highest potato cultivation in Germany (Suettdorf 53°00’39.7″N, 10°41’26.3″E; Borg 53°00’34.6″N, 10°44’33.0″E; 53°00’35.3″N, 10°44’30.2″E). Field sites were fallows next to potato growing areas and selected based on the occurrence of wireworms to ensure high pest infestation. Four weeks before the start of the experiments, the fallows were prepared by mechanical tillage.

#### Experimental design of field trials

The effectiveness of the *Metarhizium brunneum* formulations in the field was studied during three years (Suettdorf 2018, Borg 2019 and 2020) using a randomised complete block design with four replicates and an untreated control for each block. The tested treatments with the respective application rates are shown in Table 2. Individual plot size was four 10 m rows (30 m²) and edge effects were avoided by assessing potatoes from the middle two rows. Potatoes were cultivated according to agricultural practice and growth stages were determined with the BBCH code of Hack et al. (1993). The potato (*Solanum tuberosum*) variety Princess (Solana GmbH & Co.KG, Hamburg, Germany) was used in all trials with a planting density of 25 dt ha^-1^ and 0.33 m planting distance within the row. Field trials were carried out in cooperation with the Lower Saxony Chamber of Agriculture.

**Table 1:** *Metarhizium brunneum* formulations, application rates, abbreviations and seed dressing in field trials near Uelzen from 2018 to 2020.

| Year | Treatment | Application rate | Abbr. | Seed dressing |
| --- | --- | --- | --- | --- |
| 2018 | ATTRACAP® (Reference) | 30 kg ha<sup>-1</sup> | A30 | Moncut® |
|  | Granule | 30 kg ha<sup>-1</sup> | G30 | Moncut® |
|  | Dry Product | 1x10<sup>11</sup> spores ha<sup>-1</sup> | DP1 | Moncut® |
|  | Granule + Dry Product | 30 kg ha<sup>-1</sup> + 1x10<sup>11</sup> spores ha<sup>-1</sup> | G30+DP1 | Moncut® |
|  | Granule + Dry Product | 15 kg ha<sup>-1</sup> + 0.5x10<sup>11</sup> spores ha<sup>-1</sup> | G15+DP0.5 | Moncut® |
| 2019 | ATTRACAP® (Reference) | 30 kg ha<sup>-1</sup> | A30 | Moncut® |
|  | Granule | 30 kg ha<sup>-1</sup> | G30 | Moncut® |
|  | Granule | 60 kg ha<sup>-1</sup> | G60 | Moncut® |
|  | Dry Product | 1x10<sup>11</sup> spores ha<sup>-1</sup> | DP1 | Moncut® |
|  | Dry Product | 2x10<sup>11</sup> spores ha<sup>-1</sup> | DP2 | Moncut® |
|  | Granule + Dry Product | 30 kg ha<sup>-1</sup> + 1x10<sup>11</sup> spores ha<sup>-1</sup> | G30+DP1 | Moncut® |
| 2020 | ATTRACAP® (Reference) | 30 kg ha<sup>-1</sup> | G30 | Moncut® |
|  | Granule | 30 kg ha<sup>-1</sup> | G30 | Moncut® |
|  | Granule | 60 kg ha<sup>-1</sup> | G60 | Moncut® |
|  | Dry Product | 1x10<sup>11</sup> spores ha<sup>-1</sup> | DP1 | Moncut® |
|  | Granule + Dry Product | 30 kg ha<sup>-1</sup> + 1x10<sup>11</sup> spores ha<sup>-1</sup> | G30+DP1 | Moncut® |

**Table 2:**
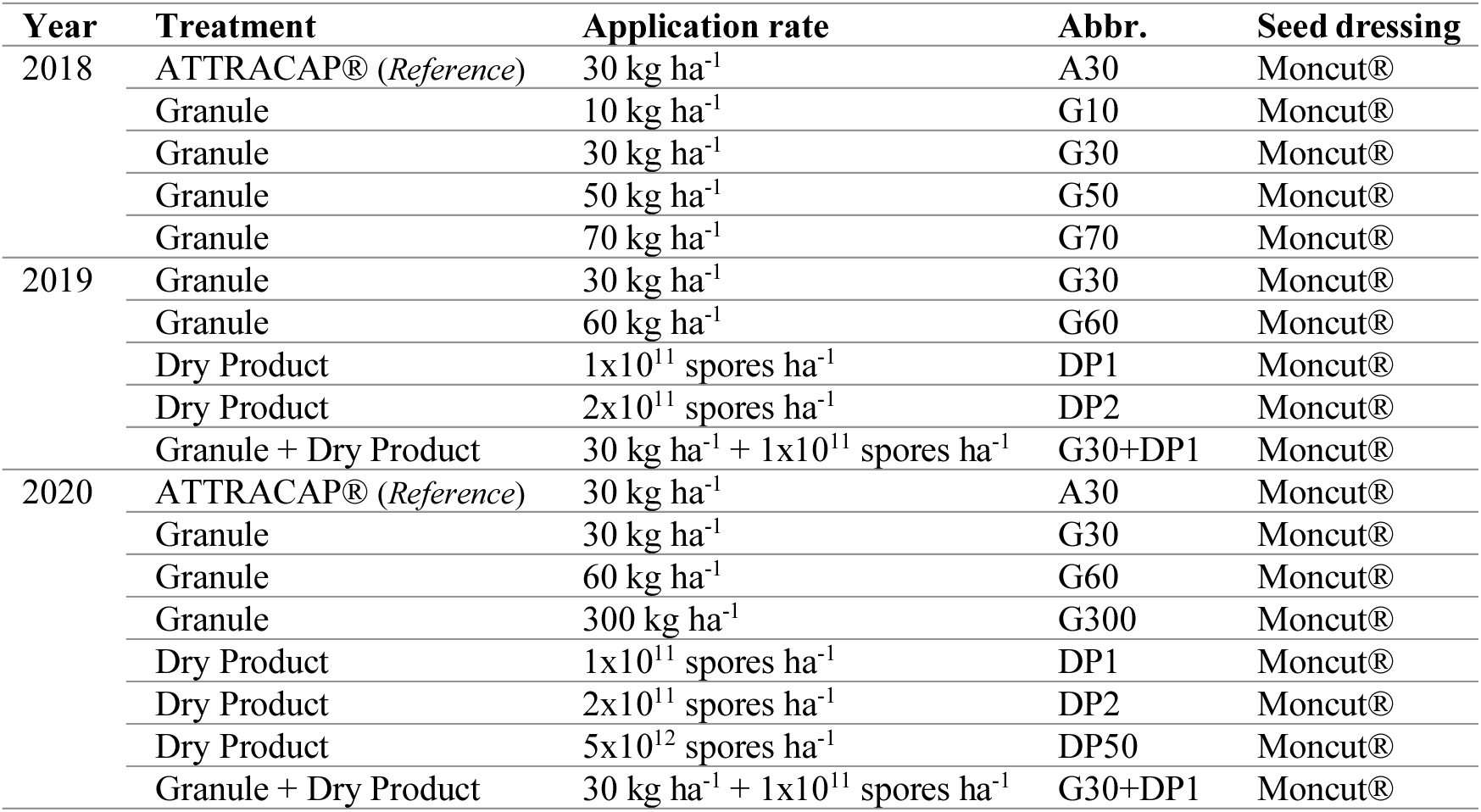
*Metarhizium brunneum* formulations, application rates, abbreviations and seed dressing in greenhouse experiments from 2018 to 2020.

#### Application of Metarhizium brunneum formulations in the field

The AgriMet-Granule and ATTRACAP® were applied during potato planting with the AgroDos® 12-volt spreader (LEHNER Maschinenbau GmbH, Westerstetten, Germany) connected to a potato planting machine. The distributor triangle of the spreader was attached behind the planting machine blade to applicate in the furrow under the potato. To ensure protection against phytopathogenic fungi the potato seed dressing Moncut® 460 SC (Belchim Crop Protecion Deutschland GmbH, Burgdorf, Germany) was applied with an application rate of 0.2 L t^-1^ in 80 L ha^-1^ with the hollow cone nozzle ALBUZ® ATR brown (agrotop GmbH, Obertraubling, Germany). For the application of the AgriMet-Dry Product the corresponding amount was dissolved in tap water and a tank mixture with Moncut® 460 SC was prepared. The activity of the *Metarhizium brunneum* isolate JKI-BI-1450 in combination with Moncut® 460 SC was confirmed in a laboratory test system (Bernhardt et al. 2019).

#### Sampling of wireworms, CFUs and soil temperature/moisture/properties

Wireworms were sampled at four different dates to investigate the predominant species and field distribution. Eight samples per plot were taken randomly at 15 cm depth with a cylindrical (10 cm diameter) auger. Wireworms were extracted by hand and determined to species level based on their morphology following Klausnitzer (1994) and Cocquempot et al. (1999).

To investigate the abundance and persistence of *Metarhizium* spp., the colony-forming units (CFUs) were analysed 0, 27, 48, 83 and 111 days after application. Five samples per plot were randomly taken in the middle of the potato ridge at 5-15 cm depth with a cylindrical (2.5 cm diameter) auger (Baumann Saatzuchtbedarf, Waldenburg, Germany). Soil samples were transferred in plastic bags (ISTAD, IKEA) and stored at 5 °C until processing. For the isolation of vital fungal cells, 20 g of each sieved (1 mm mesh) soil sample was transferred into a 300 ml flask (Schott AG, Mainz, Germany) and 100 ml 0.1 % Tween80® (Carl Roth GmbH & Co. KG, Karlsruhe, Germany) was added. After 2 h on a shaker (CERTOMAT® R, B. Braun Melsungen AG, Melsungen, Germany) at room temperature, 100 µl supernatant was spread on each of three Petri dishes with semi-selective media (Strasser et al. 1996). The product Syllit® (Arysta LifeScience Germany GmbH, Düsseldorf, Germany) was used for the addition of Dodine. The plates were incubated at 25 °C in the dark. After 14 days, the CFUs of *Metarhizium* spp. were identified by morphological criteria and counted to calculate the CFUs g^-1^ soil (Zimmermann 2007).

Soil temperature and soil moisture were measured every hour over the entire trial period placing three HOBO Micro Station H21-USB datalogger (Onset Computer Cooperation, Bourne, USA) randomly on the field. Each datalogger was equipped with a 12-Bit Temperature Smart Sensor S-TMB-MOxx (Onset Computer Cooperation, Bourne, USA) and a Soil Moisture Smart Sensor S-SMx-M005 (Onset Computer Cooperation, Bourne, USA) which were installed in the potato ridge at seed potato level (∼ 15 cm depth). The data were collected with the program HOBOware version 3.7.12 (Onset Computer Cooperation, Bourne, USA) and processed in Excel (Microsoft Office 2016) to calculate the daily mean, minimum and maximum temperature (°C) and water content (m³ m³^-1^).

The analysis of the soil properties of the three field sites was carried out by the LUFA Nord-West (Oldenburg, Germany). For this purpose, twenty samples were taken evenly distributed over the respective field site using a cylindrical (2.5 cm diameter) auger (Baumann Saatzuchtbedarf, Waldenbrug, Germany) at a depth of 30 cm. Samples were collected in a bucket and 500 g of composite samples of each field site were sent to the LUFA Nord-West in sealed bags for the analysis of copper (Cu), the humus content, the carbon-nitrogen ratio (C/N), the pH-value and the soil composition.

### Effectiveness of the *Metarhizium brunneum* formulations in the greenhouse

#### Experimental design in the greenhouse

A greenhouse experiment was carried out over three years (2018-2020) to test the effectiveness of the *Metarhizium brunneum* formulations under standardised conditions. Pots (625 cm² surface, 28 cm height) were filled with 7.5 kg of a soil (Einheitserde Classic Pikiererde CL P, Gebrüder Patzer GmbH & Co. KG, Sinntal, Germany) and sand mixture (5:1). Five *Agriotes obscurus* larvae, one seed potato (*Solanum tuberosum*) of the variety Princess (Kartoffel-Müller, Nersingen, Germany) and the respective treatment were added for one replicate. To adapt the distribution of the potato seed dressing Moncut® 460 SC with an application rate of 0.2 L t^-1^ in 80 L ha^-1^, the same nozzle as used in the field was attached to a pressure vessel. Each bucket was treated with a pressure of 6 bar for 3 sec on the soil surface at a height of 12 cm. As in the field, the corresponding amount of AgriMet-Dry Product was dissolved in tap water and applied in combination with Moncut® 460 SC. The AgriMet-Granule and ATTRACAP® were applied by hand at the same height as the AgriMet-Dry Product. Based on the research of Eckhard Immenroth regarding to the distribution of the AgriMet-Granule in the potato ridge (personal communication, 2019, JKI, Institute for Application Techniques in Plant Protection, Braunschweig, Germany), the application rate in the field could be transferred to the pots. The calculation of the application rate for the AgriMet-Dry Product was based on the pot surface. After the application, one plant potato was placed in the middle of the bucket and filled up with 12 cm of the soil-sand mixture. The pots were set up in a randomised complete block design with six replicates and stored at 20 °C and 60 % RH from April to July in the greenhouse with a daily exposure of 16 h light and 8 h darkness. The tested treatments are shown in Table 3. After planting, no plant protection product was used and the pots were watered as needed.

**Table 3:** *Metarhizium brunneum* formulations, application rates and abbreviations in the laboratory bioassay.

| Treatment | Application rate | Abbr. |
| --- | --- | --- |
| Granule | 30 kg ha<sup>-1</sup> | G30 |
| Granule | 300 kg ha<sup>-1</sup> | G300 |
| Granule | 3000 kg ha<sup>-1</sup> | G3000 |
| Granule | 15000 kg ha<sup>-1</sup> | G15000 |
| Dry Product | 1x10<sup>11</sup> spores ha<sup>-1</sup> | S1 |
| Dry Product | 1x10<sup>12</sup> spores ha<sup>-1</sup> | S10 |
| Dry Product | 5x10<sup>12</sup> spores ha<sup>-1</sup> | S50 |
| Dry Product | 1x10<sup>13</sup> spores ha<sup>-1</sup> | S100 |

#### Assessment of wireworm survival in the greenhouse

At the end of the experiment, mortality of the released *Agriotes obscurus* larvae was examined by thoroughly searching the substrate of each pot. Wireworms that were not recaptured were assessed as dead, as an escape could be excluded.

### Effectiveness of the *Metarhizium brunneum* formulations in the laboratory

#### Bioassay for the LC_50_ and LT_50_ determination

The lethal concentration (LC_50_) of the AgriMet-Granule and the AgriMet-Dry Product against *Agriotes obscurus*, *A. sputator* and *A. lineatus* was examined in a laboratory experiment using four different application rates shown in Table 4, in comparison with an untreated control.

**Table 4:** Quality control of the *Metarhizium brunneum* formulations tested. Mean outgrowth rate in percent of the AgriMet-Granule and ATTRACAP® and mean number of colony-forming unit’s (CFUs) of *M. brunneum* per g AgriMet-Dry Product for the application in field, greenhouse and laboratory experiments in 2018 to 2020. The standart deviation is shown as ±SD.

| Year | Experiment | AgriMet-Granule |  | ATTRACAP® |  | AgriMet-Dry Product |  |
| --- | --- | --- | --- | --- | --- | --- | --- |
|  |  | Outgrowth rate in % |  |  |  | CFUs g<sup>-1</sup> |  |
|  |  | mean | ±SD | mean | ±SD | mean | ±SD |
| 2018 | Field trial | 79 | 6.63 | 100 | 0.00 | 6.3x10<sup>8</sup> | 5.46x10<sup>7</sup> |
|  | Greenhouse | 81 | 7.35 | 99 | 2.0 | / | / |
| 2019 | Field trial | 78 | 5.10 | 98 | 2.5 | 8.9x10<sup>8</sup> | 6.4x10<sup>7</sup> |
|  | Greenhouse | 69 | 3.74 | 99 | 2.0 | 8.3x10<sup>8</sup> | 6.0x10<sup>7</sup> |
|  | Laboratory | 70 | 3.16 | / | / | 5.6x10<sup>8</sup> | 8.5x10<sup>7</sup> |
| 2020 | Field trial | 80 | 3.16 | 97 | 4.0 | 7.2x10<sup>8</sup> | 6.2x10<sup>7</sup> |
|  | Greenhouse | 82 | 4.00 | 98 | 2.5 | 9.6x10<sup>8</sup> | 7.7x10<sup>7</sup> |

Small plastic cans (50 ml, OPTIMAX Packaging GmbH & Co KG, Norderstedt, Germany) were filled with 35 g of a 5:1 mixture of soil (Einheitserde Classic Pikiererde CL P, Gebrüder Patzer GmbH & Co. KG, Sinntal, Germany) and sand. After the appropriate amount of AgriMet-Granule was mixed in evenly or the dissolved AgriMet-Dry Product was applied with a pipette respectively, one wireworm was added. As in the greenhouse, the application rate was calculated based on previous research and the surface of the cans used. Here, no potato seed dressing was applied. Plastic cans were maintained under controlled conditions (25 °C; 60 RH) for 105 days and wireworms were fed with *Triticum aestivum* seeds (Cultivar: Primus, Deutsche Saatgutveredelung AG, Lippstadt, Germany) to avoid starving. Larval mortality caused by *M. brunneum* was assessed two times a week by the examination of fungal outgrowth of the cadaver to determine the lethal time of 50 % mortality (LT_50_). Each treatment contained six wireworms of the respective wireworm species and the bioassay was repeated five times time independent. Thus, a total of 810 larvae were tested.

### Statistical analyses

Statistical analyses were carried out using the software R Studio (Version 1.4.1106) (RStudio Team 2020). Data are presented as the arithmetic mean (mean) and standard deviation (±SD) or adjusted mean (adjusted mean) and standard error (±SE), depending on whether a model was used for calculation. To meet the ANOVA’s underlying assumptions of normal distribution and variance homogeneity, residuals of the respective model were visually inspected. Normal distribution was checked with the QQ-Plot (sample quantile-theoretical quantile) and variance homogeneity with the Residuals-Prediction-Plot. Selection of the best-fitted model was based on the Akaike Information Criterion (AIC) (Burnham & Anderson 2002) after backward elimination of the full model. Unless otherwise noted, the post hoc Tukey HSD test (*α* = 0.05) was performed with the R package *emmeans* (Lenth et al. 2018) to examine differences between the respective factor levels in the context on an ANOVA.

The influence of the temperature and field soil (explanatory variables) on the fungal growth on the surface of the AgriMet-Granule (target variable) was determined by comparing the number of conidia per grain within a Linear Model (LM). Data were log transformed to ensure normal distribution and variance homogeneity.

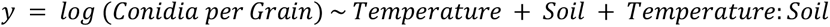

The effect of the tested *Metarhizium* formulations (explanatory variable) on the wireworm tuber damage (target variable) was analysed with a Generalized Linear Model (GLM, binominal distribution) comparing the proportion of undamaged potatoes of each plot (field) or pot (greenhouse). Global effects were assessed by performing an analysis of deviance.

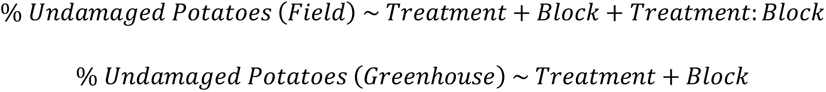

Percent treatment effectiveness was calculated relative to the untreated control for each block using Abbott’s formula (Abbott 1925).

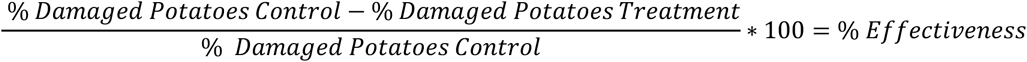

The abundance and persistence of *Metarhizium* spp. in the soil (target variable) after application of the different *M. brunneum* formulations (explanatory variable) was described as number of CFU per g soil. After square-root transformation of the CFU data, normal distribution and variance homogeneity were confirmed. Subsequent analysis of treatment effect was performed for each experimental year by setting up a Linear Mixed Model (LMM) with the *lme4* R package (Bates et al. 2015) and running an analysis of variance. A random effect was included to account for the subsamples within each plot.

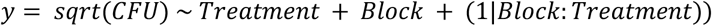

Kaplan-Meier-Analysis was used to determine the survival probability of the tested *A. obscurus*, *A. sputator* and *A. lineatus* larvae over time after incubation with different application rates of the AgriMet-Granule and AgriMet-Dry Product. Survival curves over time were created using the *survival* R package (Therneau 2021) and compared with the log-rank test to determine global differences. To detect significant differences between the application rates within a wireworm species a pairwise comparison of survival curves based on the Bonferroni method was carried out by using the *survminer* R package (Kassambara et al. 2021). To determine the time at which 50 % of the tested individuals died due to the fungal treatment (LT_50_), the “surv_median()” command of the *survival* R package was used. Mycosis of wireworms was recorded as an “event”. If no event was observed by the end of the study, the total survival time could not be accurately determined and was censored. To estimate the LC_50_ values of the AgriMet formulations, a probit analysis was performed using the *ecotox* R package (Hlina et al. 2019)

The density of wireworms on the respective field site was plotted using the R package *desplot* (Wright 2020). Recorded soil temperature, soil water content and precipitation during the field trials were visualised using Microsoft Excel (Version 2016). All other graphs were created with the R packages *ggplot2* (Wickham 2016), *ggpubr* (Kassambara 2020), *RColorBrewer* (Neuwirth 2014), and *multcompView* (Graves et al. 2019).

## Results

### Quality control of the *Metarhizium brunneum* formulations

The functionality of the different *Metarhizium brunneum* formulations was examined to ensure comparability and verify the manufactures specifications. The manufactures of the AgriMet formulations determined an outgrowth rate for the Granule between 81-100 % and 1.25×10^9^ CFU g^-1^ for the AgriMet-Dry Product. Since ATTRACAP® is commercially available, a 100 % outgrowth rate was expected. Table 5 shows that the outgrowth rate of the used AgriMet-Granule fluctuated and ranged between 69-82 % for all trials, whereas ATTRACAP® achieved an outgrowth rate between 97-100 %. The range of the AgriMet-Dry Product was between 5.6×10^8^-9.6 ×10^8^ CFUs g^-1^.

**Table 5:** Effectiveness of the *Metarhizium brunneum* formulations as the percentage of reduced potato damage compared to the untreated control during field trials in 2018 to 2020 (A30 = ATTRACAP® 30 kg h^-1^; G30 = AgriMet-Granule 30 kg ha^-1^; G60 = AgriMet-Granule 60 kg ha^-1^; DP0.5 = AgriMet-Dry Product 0.5×10^11^ conidia ha^-1^; DP1= AgriMet-Dry Product 1×10^11^ conidia ha^-1^; DP2 = AgriMet-Dry Product 2×10^11^ conidia ha^-1^). Mean and standart deviation (±SD) per year was calculated from the effectiveness for each treatment within a block according to Abbott (1925).

| Treatment | Effectiveness [%] |  |  |  |  |  |
| --- | --- | --- | --- | --- | --- | --- |
|  | 2018 |  | 2019 |  | 2020 |  |
|  | mean | ±SD | mean | ±SD | mean | ±SD |
| A30 | 11.3 | 7.2 | 9.0 | 12.2 | 2.1 | 2.9 |
| G30 | 2.0 | 1.4 | -2.9 | 23.9 | 6.6 | 5.2 |
| G60 | / | / | 11.4 | 24.1 | 8.7 | 13.9 |
| DP1 | 11.9 | 5.4 | 4.4 | 17.1 | 6.3 | 9.2 |
| DP2 | / | / | -4.9 | 36.9 | 3.1 | 3.2 |
| G30_DP1 | 9.6 | 10.1 | 13.6 | 16.3 | 1.0 | 1.8 |
| G15_DP0.5 | 3.4 | 2.8 | / | / | / | / |

The quality control of the AgriMet-Granule in field soil revealed that the initial conidia concentration of the *M. brunneum* isolate JKI-BI-1450 coated on the surface of the tested batch was 5.89×10^4^ conidia grain^-1^ (±SD 5.05×10^3^ conidia grain^-1^). The sporulation after 14 days of incubation was significant influenced by the tested temperatures and respective substrate (ANOVA, *p* < 0.0001). Figure 8 shows that notable sporulation of the AgriMet-Granule was only visible on the sterile control substrate 1.5 % water agar. The incubation temperature of 10 °C increased the conidia concentration only slightly to 1×10^5^ conidia grain^-1^. In comparison, a significant increase of the *M. brunneum* concentration was determined at 15 °C with 5.6×10^5^ conidia grain^-1^ (Tukey HSD test, *p* < 0.0001). After an exponential increase to 1.1×10^7^ conidia grain^-1^ at 20 °C (15-20°C: Tukey HSD test, *p* < 0.0001), sporulation reached its maximum at 25 °C with 2.3×10^7^ conidia grain^-1^. In contrast, the incubation of the AgriMet-Granule on the soil from Suettdorf or Borg led to no significant increase in conidia formation up to a temperature of 20 °C. Only the incubation at 25 °C resulted in a significant but small increase to 3.9×10^5^ (Suettdorf) and 7.1×10^5^ (Borg) conidia grain^-1^. Numerous grains incubated in the respective soil were found to be colonised by other fungi and bacteria.

**Figure 1:**
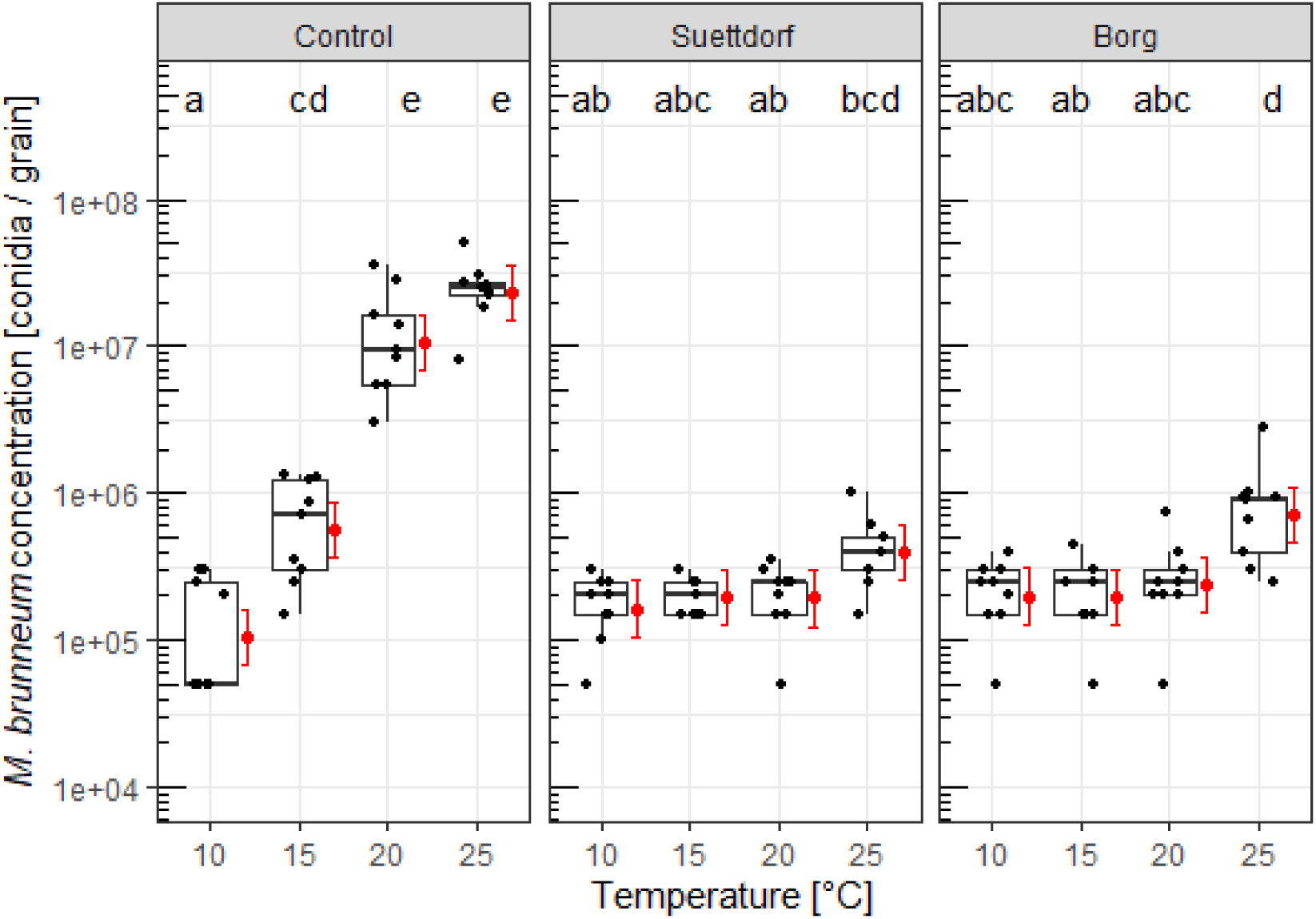
*Metarhizium brunneum* concentration (conidia grain^-1^) on the AgriMet-Granule after incubation at 10 °C, 15 °C, 20 °C and 25 °C on 1.5 % water agar (Control) or field soil from Suettdorf and Borg for 14 days. Black dots per temperature and substrate represent the mean number of conida grain^-1^ out of ten grains per Petri Dish. Boxplots consist of the median and the 25 % and 75 % quantile. Adjusted mean of conidia grain^-1^ and 95 % confidence interval of the replicates per treatment are illustrated in red and treatments with the same letters are not significantly different. LM (y = log(Conidia per Grain) ∼ Temperature + Soil + Temperature:Soil), Tukey HSD test (*α* = 0.05).

### Effectiveness of the *Metarhizium brunneum* formulations in the field

#### Assessment of wireworm potato damage

The effectiveness of the tested *Metarhizium brunneum* formulations in field trials over three years was assessed by the proportion of undamaged potatoes after harvest and the results are shown in Figure 9. A very high infestation of wireworms in each year was reflected by very small proportions of undamaged potatoes in the untreated controls that ranged between 2.3 % (2018), 13.1 % (2019) and 0.3 % (2020). The tuber damage was significantly reduced by the application of individual *M. brunneum* formulations in each year with strong block effects (Treatment: ANOVA, *p* < 0.0001, Block: ANOVA, *p* < 0.0001). However, the effectiveness of the treatments was very low and neither the reference product ATTRACAP® nor the AgriMet formulations had a constant effect over all years. In 2018, the application of ATTRACAP® reduced the tuber damage significantly to a proportion of 12 % undamaged potatoes compared to the untreated control (Tukey HSD test, *p* < 0.001). A similar effect was achieved by the combination of the AgriMet-Granule with 30 kg ha^-1^ plus the AgriMet-Dry Product with 1×10^11^ conidia ha^-1^ (10 %) as well as with the AgriMet-Dry Product with 1×10^11^ conidia ha^-1^ individually (9 %), with no difference between treatments (A30-G30_DP1: Tukey HSD test, *p* = 0.99; A30-DP1: Tukey HSD test, *p* = 0.88, G30_DP1-DP1: Tukey HSD test, *p* = 0.99). In 2019, the increased application rate of the AgriMet-Granule to 60 kg ha^-1^ resulted in the highest proportion of undamaged potatoes of 27 % with a significant difference of the untreated control (Tukey HSD test, *p* < 0.0001). As in the previous year, the potato damage was comparable within the treatments ATTRACAP® (23 %) and the combination of the AgriMet-Granule with 30 kg ha^-1^ plus AgriMet-Dry Product with 1×10^11^ conidia ha^-1^ (26 %) (G60-A30: Tukey HSD test, *p* = 0.78; G60-G30_DP1: Tukey HSD test, *p* = 0.99; A30-G30_DP1: Tukey HSD test, *p* = 0.96). Although the AgriMet-Dry Product with 1×10^11^ conidia ha^-1^ individually increased the proportion of undamaged potatoes, there was no significant difference to the control (Tukey HSD test, *p* = 0.09). In addition, a strong block effect was obvious with a large scattering within the treatments (ANOVA, *p* < 0.0001). The majority of plots in block A showed the lowest proportion of undamaged potatoes regardless of the treatment (1-21 %), whereas the proportion in plots from block B was highest (25-56 %). In 2020, wireworm damage was extremely high with 0.3 % undamaged potatoes in the untreated control. The application of ATTRACAP® resulted in 1 % undamaged potatoes with no significant difference to the control (Tukey HSD test, *p* = 0.40). The *Metarhizium* formulation with the best effectiveness was the AgriMet-Granule with 60 kg ha^-1^ and a proportion of 4 % undamaged potatoes (G60-C: Tukey HSD test, *p* = 0.0003; G60-A30: Tukey HSD test, *p* = 0.0049). However, the scattering within this treatment was relatively wide and ranged from 0 % (Block A) to 34 % (Block C) undamaged potatoes. A similar scattering between 0 % (Block C) and 23 % (Block D) of the AgriMet-Dry Product with 1×10^11^ conidia ha^-1^ resulted in 2.8 % undamaged potatoes (DP1-G60: Tukey HSD test, *p* = 0.82). The AgriMet-Dry Product as well as the AgriMet-Granule with 30 kg ha^-1^ (3.1 %) led a small but significant increase in the proportion of undamaged potatoes compared to the control with no difference between the treatments (DP1-C: Tukey HSD test, *p* = 0.0045, G30-C: Tukey HSD test, *p* = 0.0019, DP1-G30: Tukey HSD test, *p* = 0.99). The application of the AgriMet-Dry Product with 2×10^11^ conidia ha^-1^ as well as the combination of the AgriMet-Granule with 15 kg ha^-1^ plus the AgriMet-Dry Product with 0.5×10^11^ conidia ha^-1^ did never reduce the tuber damage significantly.

**Figure 2:**
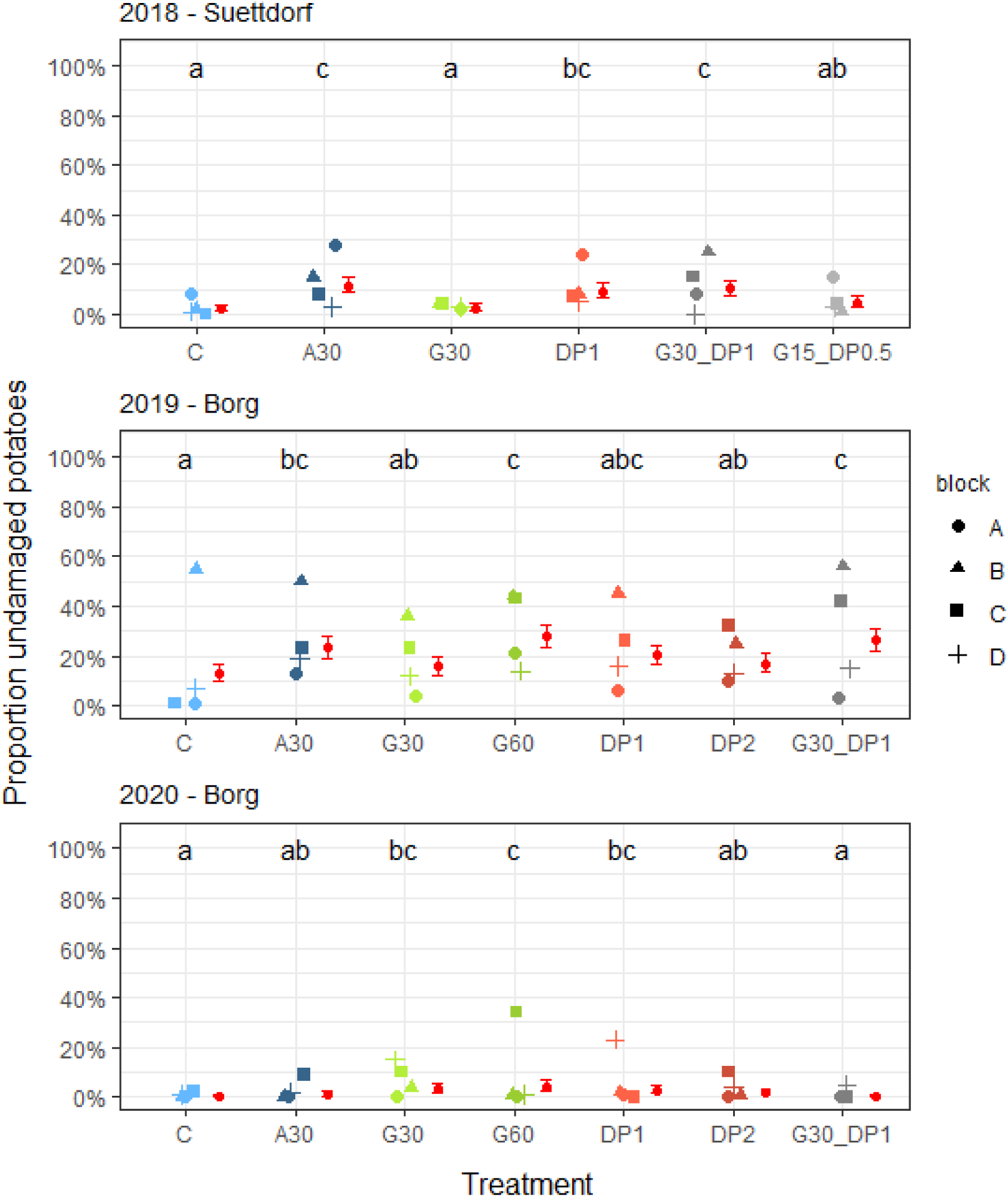
Proportion of undamaged potatoes after harvest of field trials with different formulations of *Metarhizium brunneum* from 2018 to 2020 near Uelzen. The four symbols per treatment (C = untreated control; A30 = ATTRACAP® 30 kg h^-1^; G30 = AgriMet-Granule 30 kg ha^-1^; G60 = AgriMet-Granule 60 kg ha^-1^; DP0.5 = AgriMet-Dry Product 0.5×10^11^ conidia ha^-1^; DP1= AgriMet-Dry Product 1×10^11^ conidia ha^-1^; DP2 = AgriMet-Dry Product 2×10^11^ conidia ha^-1^) represent the percentage of undamaged potatoes out of 100 randomly sampled tubers per plot within each block in a randomized complete block design. Adjusted mean and 95 % confidence interval of the four plots per treatment are illustrated in red and treatments with the same letters within a year are not significantly different. GLM (y = Percentage Undamaged Potatoes ∼ Treatment + Block + Treatment:Block, family=binominal), Tukey HSD test (*α* = 0.05).

The effectiveness of the *M. brunneum* formulations compared to the untreated control according to Abbott (1925) is shown in Table 6 and underlines the statistically analysed low tuber damage reduction. The reference product ATTRACAP® (30 kg ha^-1^) reached a three-year-average effectiveness of 7.5 %, whereas the same application rate of AgriMet-Granule resulted in only 1.9 %. The increased application rate of the AgriMet-Granule with 60 kg ha^-1^ led to 10.5 % effectiveness in average. Although the AgriMet-Dry Product with 1×10^11^ conidia ha^-1^ already reached 7.5 % effectiveness, the doubled application rate did not lead to any improvement. The combined application of the AgriMet-Granule (30 kg ha^-1^) and the AgriMet-Dry Product (1×10^11^ conidia ha^-1^) resulted in the highest effectiveness of 13.6 % compared to the untreated control in 2019 and an average of 8.0 % over all years.

**Table 6:** Soil properties of the three field sites in Suettdorf and Borg during the testing of the *M. brunneum* formulations from 2018 to 2020.

| Year | Field site | Soil analysis |  |  |  |  |  |  |
| --- | --- | --- | --- | --- | --- | --- | --- | --- |
|  |  | Copper<br>[mg kg<sup>-1</sup>] | Humus<br>content<br>[%] | C/N<br>ratio | pH<br>value | Soil components<br>[%] |  |  |
|  |  |  |  |  |  | clay | silt | sand |
| 2018 | Suettdorf | 3.8 | 3.2 | 12 | 5.9 | 6.1 | 13.8 | 80.1 |
| 2019 | Borg | 1.3 | 4.6 | 14 | 5.9 | 6.0 | 15.2 | 78.8 |
| 2020 | Borg | 1.1 | 4.8 | 14 | 5.9 | 6.1 | 16.0 | 77.9 |

#### Wireworm sampling during field trials

Wireworms were extracted from eight soil samples per plot at four sampling dates to determine the species composition and density over time. The results of the wireworm species composition are shown in Figure 10. The dominant species during each field trial was *Agriotes lineatus* with a share between 60-70 % of the sampled wireworms. *Agriotes obscurus* was the second dominant species in every year with a share between 25-35 %. In 2018 and 2020, a small number of the species *Hemicrepidius niger*, *Agrypnus murinus* and *Oedostethus quadripustulatus* was determined. However, the wireworms were not evenly distributed over each field site. Figure 11 shows that the agglomeration of wireworms in 2020 was highest in block A and lowest in block C. In 2019, block B was the area with the fewest wireworms, whereas the most were in block D. There was a clear difference between block A and B to C and D in 2018. Overall, the distribution of wireworms was always nested.

**Figure 3:**
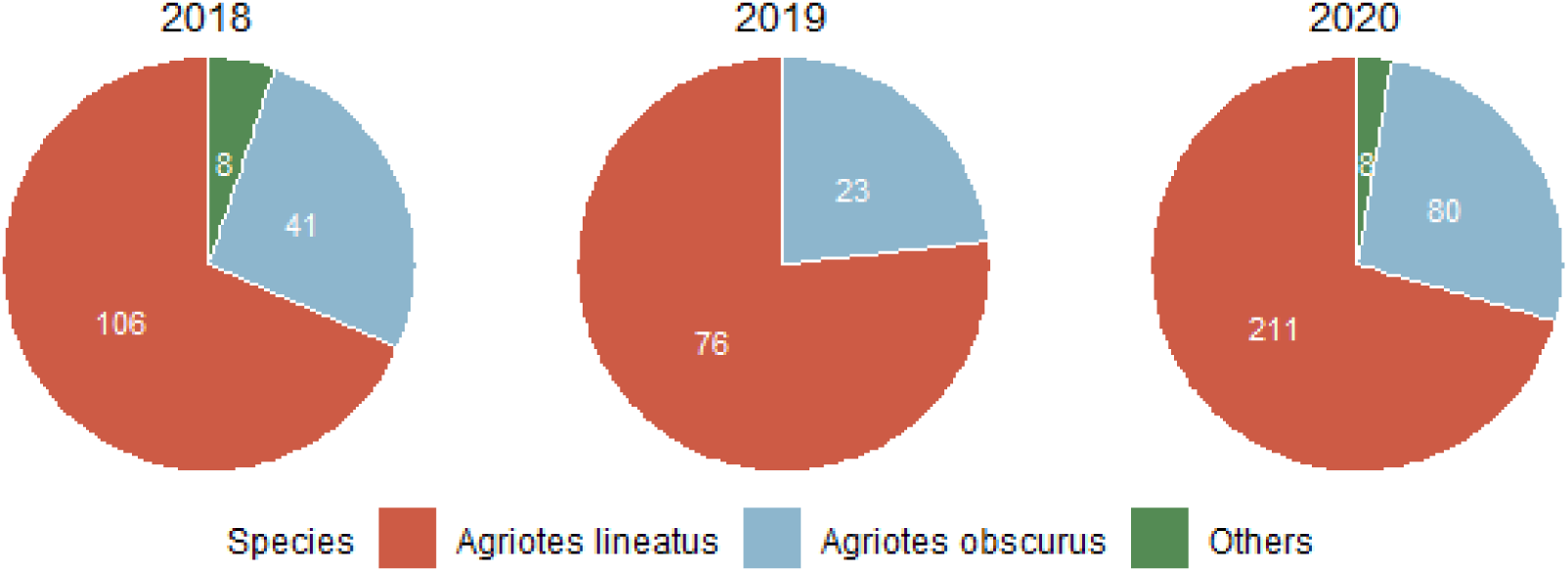
Wireworm species composition of the field trials in potatoes near Uelzen in three years. The colours represent the different wireworm species (red: *Agriotes lineatus*, blue: *A. obscurus*, green: others) and the respective total number is illustrated in white.

#### Persistence of Metarhizium spp. during field trials

The abundance and persistence of *Metarhizium* spp. was determined by the number of CFUs g^-1^ soil and the results as well as the associated statistics are shown in Table 8. There was no significant difference in the number of CFUs g^-1^ soil before the application of the formulations of *M. brunneum* (ANOVA, *p* > 0.05) in each of the three field trials. However, the initial concentration of *Metarhizium* spp. was twenty times higher in 2018 compared to 2019 and 2020. In 2018, the application of G30+DP1 was the only treatment which increased the concentration of *Metarhizium* spp. significantly 48 days after application (DAA) to a maximum of 959.9 CFUs g^-1^ soil (ANOVA, *p* = 0.03). 83 DAA the number von CFUs g^-1^ soil decreased and equalled to the initial value. In 2019, the application of G60, DP2 and G30+DP1 led to a significant increase of the CFUs g^-1^ soil repeatedly 48 DAA (ANOVA, *p* < 0.001). G60 and G30+DP1 reached an equal level of 138.8 CFUs g^-1^ soil and 154.1 CFUs g^-1^ soil, whereas DP2 achieved 87.4 CFUs g^-1^ soil. The abundance of *Metarhizium* spp. in the treatment G60 increased further to a maximum of 191.8 CFUs g^-1^ soil after 111 DAA, whereas the CFUs g^-1^ soil of the treatment G30+DP1 decreased 111 DAA. The treatment DP2 reached a maximum of 139.1 CFUs g^-1^ soil after 83 days and decreased slightly 111 DAA. In 2020, no treatment was able to increase the CFUs g^-1^ soil significant. However, the application of A30 resulted in a slight increase up to 86.1 CFUs g^-1^ soil after 83 days (ANOVA, *p* = 0.54). In addition, G60 reached 42.1 CFUs g^-1^ soil and G30+DP1 36.6 CFUs g^-1^ soil after 83 days. 111 DAA the CFUs g^-1^ soil equalled to the *Metarhizium* spp. concentration before the application of the formulations.

#### Soil temperature and water content during field trials

The soil temperature and water content in the potato ridge at a depth of ∼15 cm was measured to consider the effect of environmental factors on the effectiveness of the *Metarhizium brunneum* formulations during field trials (see Figure 12). At the beginning of the vegetation period, the temperature in the potato ridge was 16.5 °C in 2018, 9.8 °C in 2019 and 14.7 °C in 2020. The temperature increased slowly and reached 20 °C after 25 days in 2018, 40 days in 2019 and 42 days in 2020. Afterwards, the temperature was at approximately 20 °C with small fluctuations during June and July. However, the temperature in 2020 decreased after 42 days and remained below 20 °C until 101 days after the application. This resulted in the lowest average temperature of the entire vegetation period of 17.1 °C. The average temperature in 2018 was 19.4 °C and 18.7 °C in 2019. The highest temperature per day was between 23.6 °C (2018) and 25.3 °C (2019), with an absolute maximum of 29.2 °C measured in 2019 after 94 days. The increasing temperature over the vegetation period resulted in a decreasing water content in the potato ridge. The water content differed between the years at the beginning of the vegetation period and was between 0.12 m³/m³ (2018) and 0.18 m³/m³ (2019). Over time, the water content dropped due to the low total precipitation (2018: 77.6 mm; 2020: 172.7 mm) to a minimum of 0.07 m³/m³ in 2018 and 0.05 m³/m³ in 2020. However, the total precipitation in 2019 was higher (216.2 mm) and increased the falling water content from 0.07 m³/m³ to an average of 0.14 m³/m³ until harvest. In addition to the temperature and the precipitation, the water consumption of the potato plant has an influence on the water content in the potato ridge. After 60 to 80 days after planting the potato reached BBCH stage 60 and the water content in the potato ridge decreased due to tuber formation and the additional water consumption.

#### Soil properties during field trials

The soil properties of the three field sites used for testing the *M. brunneum* formulations were analysed by the LUFA Nord-West, and the results are shown in Table 7. The ratio of the soil components clay, silt and sand was nearly the same at all field sites and corresponded to that of light loamy-sandy soils. The pH-value of all soils was 5.9 and the carbon-to-nitrogen ratio (C/N ratio) varied between 12-14. The humus content was lowest in Suettdorf (2018) at 3.2 % and highest in Borg (2020) at 4.8 %. The concentration of copper ranged between 1.1-1.3 mg kg^-1^ in Borg and 3.8 mg kg^-1^ in Suettdorf. Overall, the soil properties of the different field sties were very similar and did not show any noticeable values in the analysed parameters.

**Figure 4:**
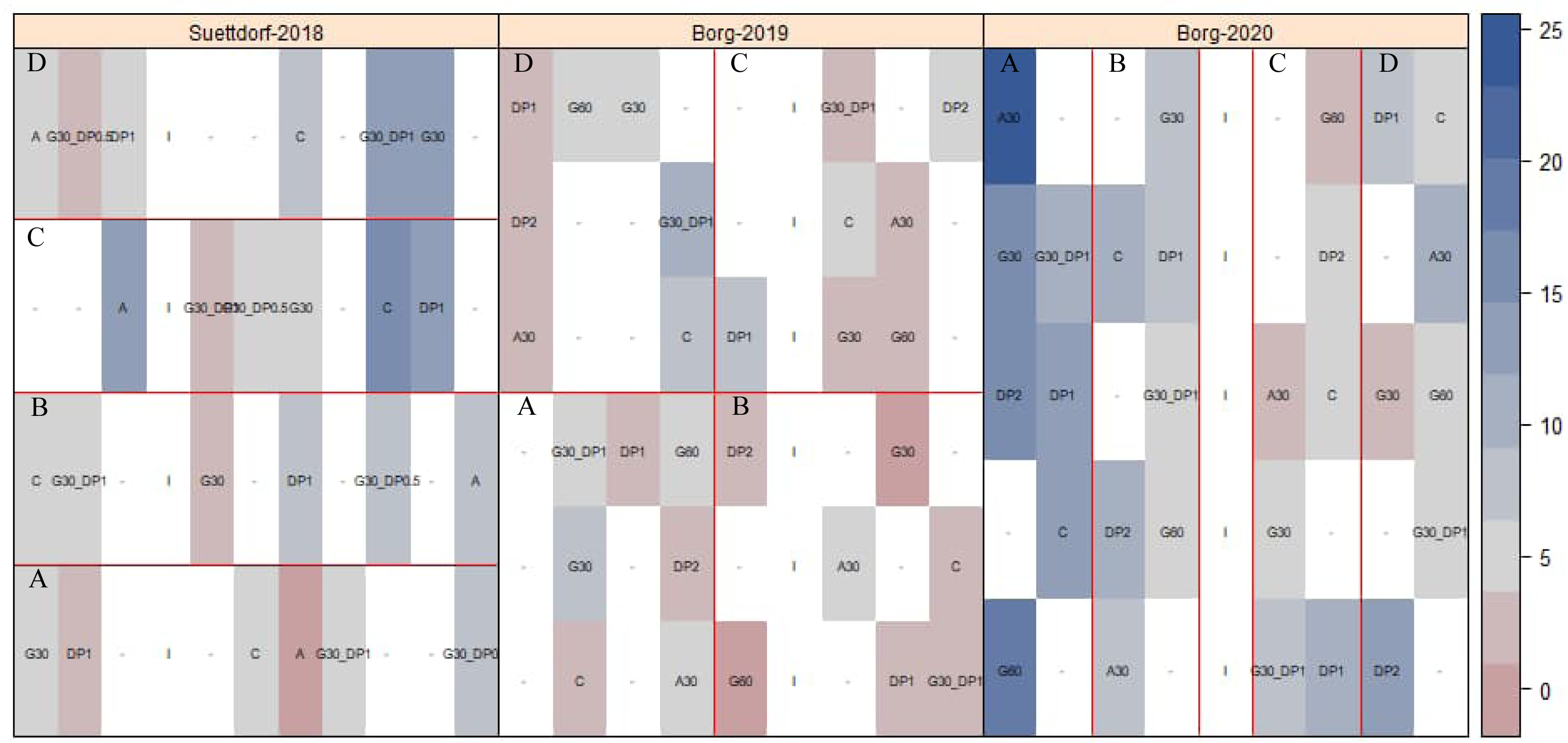
Wireworm density of the field trials in 2018 (Suettdorf), 2019 (Borg) and 2020 (Borg) based on the sum of four samplings with a cylindric auger with eigth samples per plot. The colour of each plot corresponds with the number of wireworms and is illustrated in red (no wireworms) to blue (many wireworms). The abbreviations in each plot represent the treatments (C = untreated control; A30 = ATTRACAP® 30 kg h^-1^; G30 = AgriMet-Granule 30 kg ha^-1^; G60 = AgriMet-Granule 60 kg ha^-1^; DP0.5 = AgriMet-Dry Product 0.5×10^11^ conidia ha^-1^; DP1= AgriMet-Dry Product 1×10^11^ conidia ha^-1^; DP2 = AgriMet-Dry Product 2×10^11^ conidia ha^-1^). The four blocks (A, B, C, D) per trial are separated with red lines. Plots that have not been sampled are displayed with “-“ and tramlines with “I”.

**Table 7:**
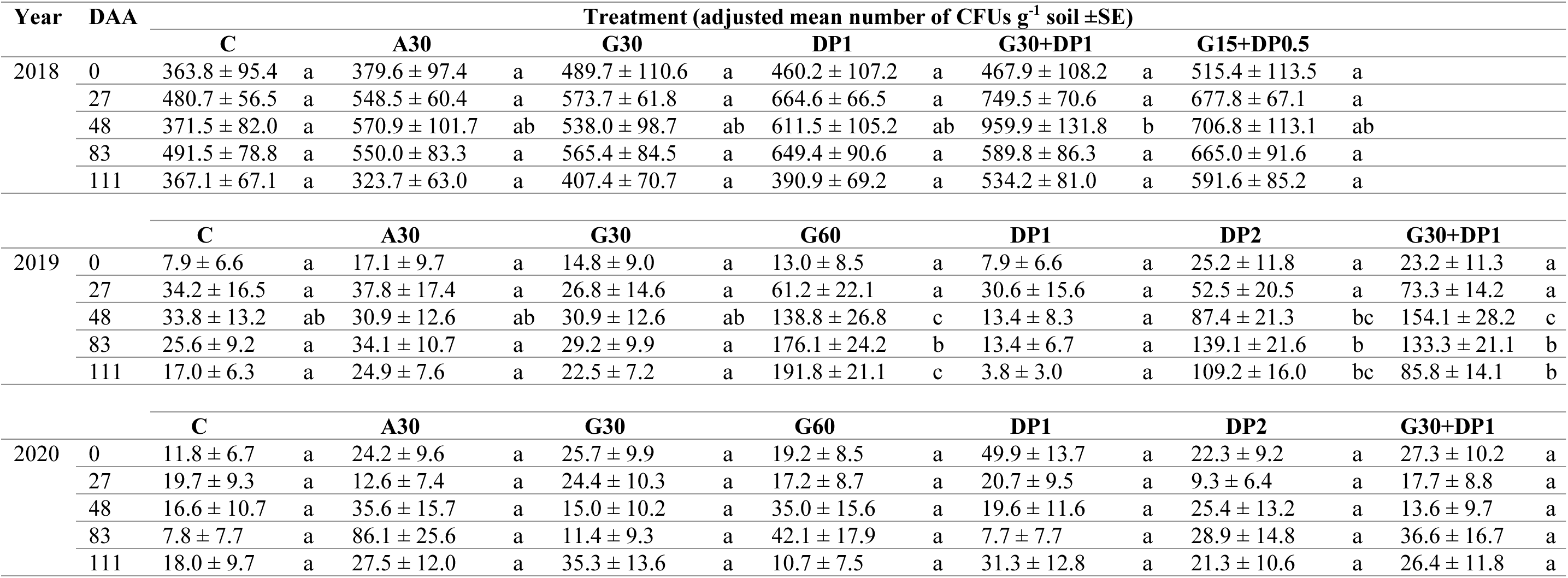
Adjusted mean number of colony-forming units (CFUs) g^-1^ soil of *Metarhizium* spp. and standard error (±SE) during field trials near Uelzen in three years before (0) and 27, 43, 83 and 111 days after the application (DAA) of different formulations von *M. brunneum* (A30 = ATTRACAP® 30 kg h^-1^; G30 = AgriMet-Granule 30 kg ha^-1^; G60 = AgriMet-Granule 60 kg ha^-1^; DP0.5 = AgriMet-Dry Product 0.5×10^11^ conidia ha^-1^; DP1= AgriMet-Dry Product 1×10^11^ conidia ha^-1^; DP2 = AgriMet-Dry Product 2×10^11^ conidia ha^-1^) and an untreated control (C). Adjusted mean values of the treatments with the same letters within a sampling date are not significantly different. LMM (y = sqrt(CFU) ∼ Treatment + Block + (1|Block:Treatment)), Tukey HSD test (*α* = 0.05).

**Figure 5:**
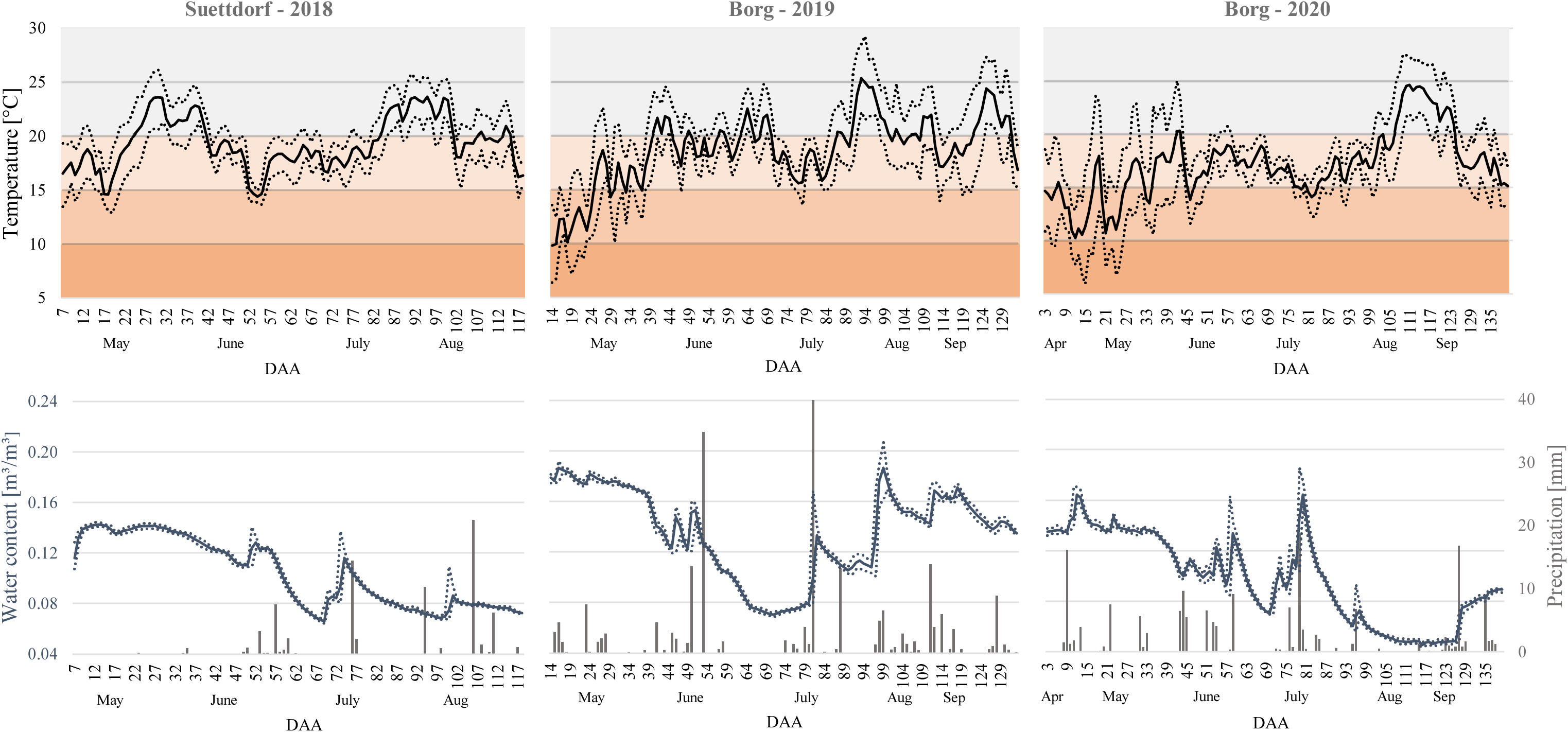
Temperature [°C] and water content [m³/m³] in the potato ridged at a depth of 15 cm after potato planting during field trials with *Metarhizium brunneum* formulations in Suettdorf (2018) and Borg (2019, 2020). Measurement was carried out by placing three HOBO-Datalogger randomly on the field site equipped with a temperature and a moisture sensor each. Daily means are presented as full lines and the maximum and minimum per day are illustrated with dotted lines. Data of the total precipitation per day was obtained from a metrological measurement station of the Deutsche Wetterdienst (DWD) in Uelzen and are shown as grey bars.

### Effectiveness of the *Metarhizium brunneum* formulations in the greenhouse

For the assessment of the appropriate application rate of the *Metarhizium brunneum* formulations AgriMet-Granule and AgriMet-Dry Product, a standardised greenhouse experiment with pots was carried out and the results are shown in Figure 13. The number of recaptured *Agriotes obscurus* larvae was not significantly influenced by any of the tested formulations. Even up to 100 times of the field application rate of the AgriMet-Granule (3000 kg ha^-1^) and the AgriMet-Dry Product (1×10^13^ conidia ha^-1^) decreased the number of wireworms only slightly compared to the control. The reference product ATTRACAP® also had no influence on the number of wireworms. Overall, the mean number of recaptured larvae was between 2.33 ± 0.36 (DP100) to 3.6 ± 0.26 (G30). However, the assessment of the tuber damage revealed differences between the treatments. The proportion of undamaged potatoes in the untreated control was 13.8 % (±SE 2.6 %) and 37.5 % (±SE 4.6 %) for the reference product ATTRACAP® (Tukey HSD test, *p* = 0.0003). The application of the AgriMet-Granule and AgriMet-Dry Product increased the proportion of undamaged potatoes significantly compared to the untreated control (ANOVA, *p* < 0.0001) with a significant impact of the application rates. While the field application rate of 30 kg ha^-1^ of the AgriMet-Granule was not able to reduce the tuber damage (Tukey HSD test, *p* = 0.98), an increased application rate of 60 kg ha^-1^ and 3000 kg ha^-1^ resulted in 42.5 % (G60) and 58.2 % (G3000) undamaged potatoes with a significant difference to the untreated control (Tukey HSD test, *p* < 0.0001). A similar result was assessed for the AgriMet-Dry Product. The application rates of 1×10^11^ conidia ha^-1^ and 2×10^11^ conidia ha^-1^ led to slightly increased proportions of 24.8 % (±SE 4.2 %) and 28.2 % (±SE 4.4 %) undamaged potatoes compared to the untreated control with no significant difference. However, 1×10^13^ conidia ha^-1^ of the AgriMet-Dry Product reduced the potato damage significant (Tukey HSD test, *p* < 0.0001) with 58.3 % (±SE 6.9 %) undamaged potatoes. The results of the high application rates of both formulations were comparable and slightly higher than the reference product ATTRACAP®. The combination of the AgriMet-Granule and the AgriMet-Dry Product resulted in 41.7 % (G30_DP1) and 55.2 % (G60_DP2) undamaged potatoes.

**Figure 6:**
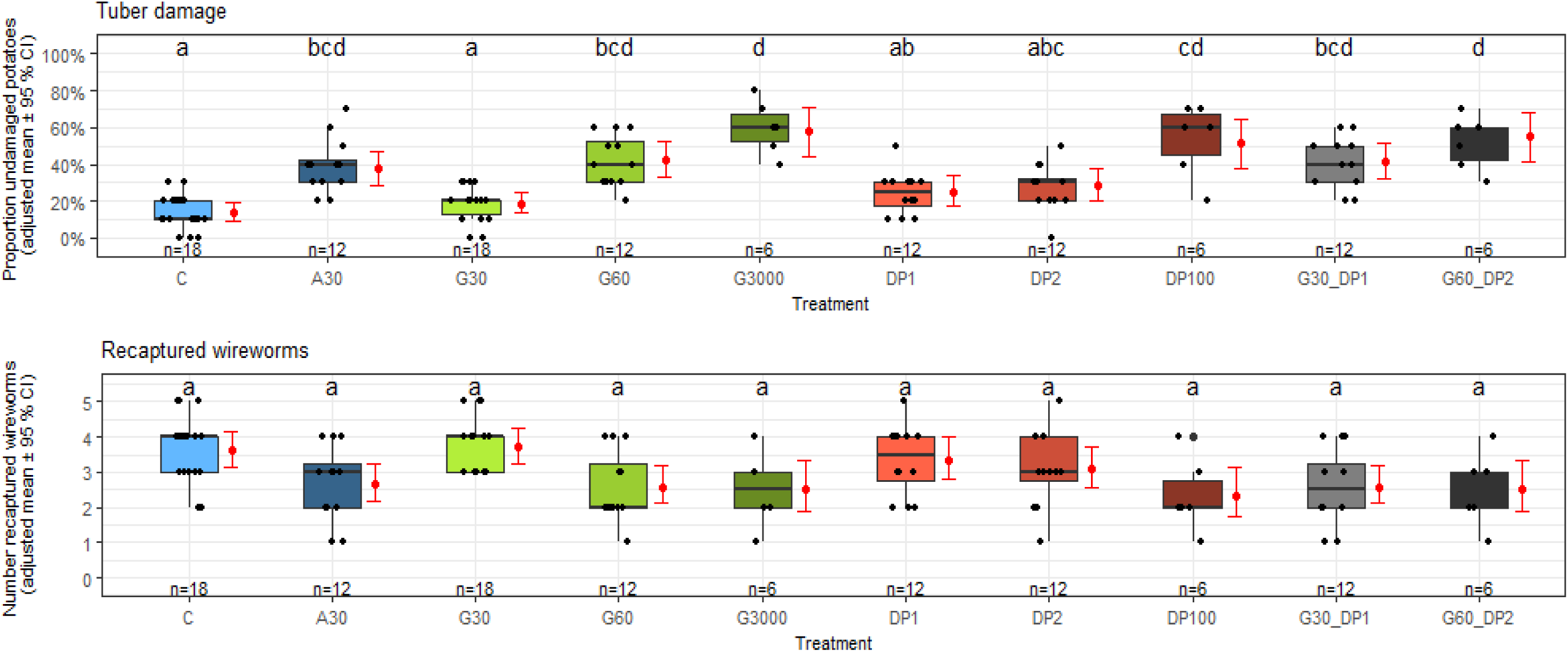
Recaptured *Agriotes obscurus* larvae and percentage of undamaged potatoes after harvest of a full potato growing season in the greenhouse experiments from 2018 to 2020. Different formulations of *Metarhizium brunneum* were applied during planting with six replicates (pots) per treatment and repetition in a randomized complete block design (A30 = ATTRACAP® 30 kg h^-1^; G30 = AgriMet-Granule 30 kg ha^-1^; G60 = AgriMet-Granule 60 kg ha^-1^; DP0.5 = AgriMet-Dry Product 0.5×10^11^ conidia ha^-1^; DP1= AgriMet-Dry Product 1×10^11^ conidia ha^-1^; DP2 = AgriMet-Dry Product 2×10^11^ conidia ha^-1^). **Top:** Dots per treatment represent the percentage of undamaged potatoes out of ten potatoes per pot. Boxplots consist of the median and the 25 % and 75 % quantile. Adjusted mean and 95 % confidence interval of n pots per treatment are illustrated in red and treatments with the same letters are not significantly different. GLM (y = Percentage Undamaged Potatoes ∼ Treatment + Year, family=binominal), Tukey HSD test *α* = (0.05). **Bottom:** Jittered boxplot of recaptured *A. obscurus* larvae per pot. Five larvae per pot were released at the beginning. Boxplots consist of the median and the 25 % and 75 % quantile. Adjusted mean of recaptured *A. obscurus* larvae and 95 % confidence interval of the replicates per treatment are illustrated in red and treatments with the same letters are not significantly different. GLM (y = Recaptured Wireworms ∼ Treatment + Year, family=poisson), Tukey HSD test (*α* = 0.05).

The effectiveness of the treatments based on the calculation according to Abbott (1925) is shown in Table 9 and indicated a dose-related reduction of tuber damage by the AgriMet formulations. The field application rates of the respective formulations resulted in an effectiveness of approximate 22 %. A 100-fold increase in the application rate of the AgriMet-Granule and AgriMet-Dry Product doubled the effectiveness to a maximum of 54.0 % and 44.1 %, respectively.

**Table 8:** Effectiveness of the *Metarhizium brunneum* formulations as the percentage of reduced potato damage compared to the untreated control during greenhouse experiments. Mean of all replicates (n) and standart deviation (±SD) was calculated from the effectiveness for each treatment within a block according to Abbott (1925).

| Treatment | Application rate | n | Effectiveness [%] |  |
| --- | --- | --- | --- | --- |
| | | | mean | ±SD |
| ATTRACAP® | 30 kg ha<sup>-1</sup> | 12 | 19.9 | 22.1 |
| AgriMet-Granule | 30 kg ha<sup>-1</sup> | 18 | 18.8 | 23.34 |
| AgriMet-Granule | 60 kg ha<sup>-1</sup> | 12 | 22.0 | 22.91 |
| AgriMet-Granule | 300 kg ha<sup>-1</sup> | 6 | 54.0 | 10.66 |
| AgriMet-Dry Product | 1x10<sup>11</sup> conidia ha<sup>-1</sup> | 12 | 24.1 | 23.43 |
| AgriMet-Dry Product | 2x10<sup>11</sup> conidia ha<sup>-1</sup> | 12 | 25.4 | 22.19 |
| AgriMet-Dry Product | 1x10<sup>13</sup> conidia ha<sup>-1</sup> | 6 | 44.1 | 23.71 |
| AgriMet-Granule + Dry Product | 30 kg ha<sup>-1</sup> + 1x10<sup>11</sup> conidia ha<sup>-1</sup> | 18 | 29.5 | 22.41 |
| AgriMet-Granule + Dry Product | 60 kg ha<sup>-1</sup> + 2x10<sup>11</sup> conidia ha<sup>-1</sup> | 6 | 42.8 | 18.29 |

**Table 9:**
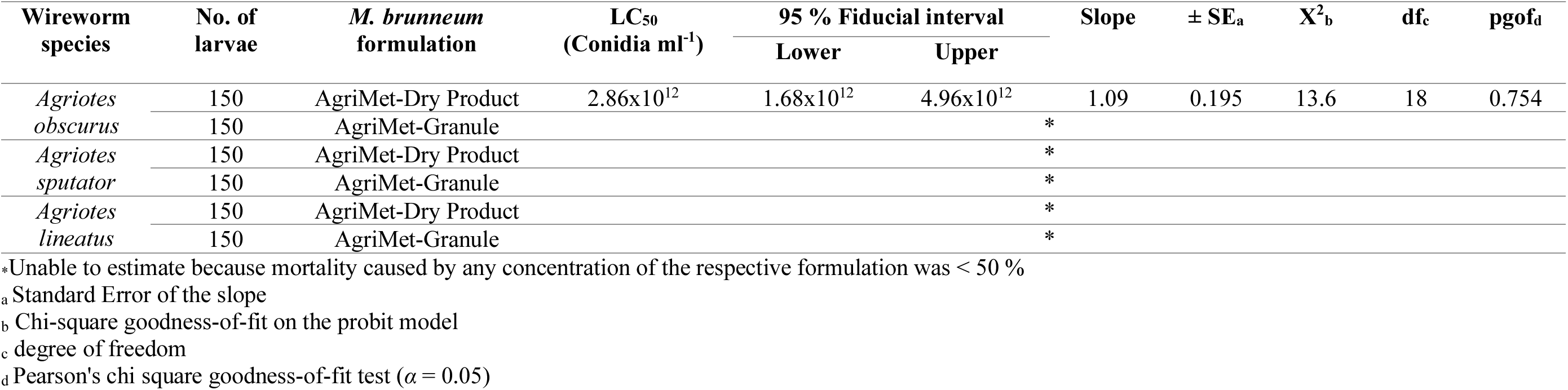
Effectiveness of the *Metarhizium brunneum* formulations AgriMet-Granule and AgriMet-Dry Product against the wireworm species *Agriotes obscurus*, *A. sputator* and *A. lineatus* after 105 days of incubation under laboratory conditions at 25 °C and darkness in small cans. Probit analysis was used to calculate the values of the lethal concentration (LC_50_) for each wireworm species exposed to the respective *M. brunneum* formulation individually.

| Wireworm species | No. of larvae | *M. brunneum* formulation | $\text{LC}_{50}$<br>(Conidia $\text{ml}^{-1}$) | 95 % Fiducial interval | | Slope | $\pm \text{SE}_a$ | $\chi^2_b$ | df<sub>c</sub> | pgof<sub>d</sub> |
| --- | --- | --- | --- | --- | --- | --- | --- | --- | --- | --- |
|  |  |  |  | Lower | Upper |  |  |  |  |  |
| *Agriotes obscurus* | 150 | AgriMet-Dry Product | $2.86 \times 10^{12}$ | $1.68 \times 10^{12}$ | $4.96 \times 10^{12}$ | 1.09 | 0.195 | 13.6 | 18 | 0.754 |
|  | 150 | AgriMet-Granule |  |  |  | * |  |  |  |  |
| *Agriotes sputator* | 150 | AgriMet-Dry Product |  |  |  | * |  |  |  |  |
|  | 150 | AgriMet-Granule |  |  |  | * |  |  |  |  |
| *Agriotes lineatus* | 150 | AgriMet-Dry Product |  |  |  | * |  |  |  |  |
|  | 150 | AgriMet-Granule |  |  |  | * |  |  |  |  |

\*Unable to estimate because mortality caused by any concentration of the respective formulation was < 50 %
<sub>a</sub> Standard Error of the slope
<sub>b</sub> Chi-square goodness-of-fit on the probit model
<sub>c</sub> degree of freedom
<sub>d</sub> Pearson's chi square goodness-of-fit test ( $\alpha = 0.05$ )

### Effectiveness of the *Metarhizium brunneum* formulations in the laboratory

A laboratory bioassay with four different application rates of the *Metarhizium brunneum* formulations AgriMet-Granule and AgriMet-Dry Product was performed to define the required amount for a reduction of the wireworm species *Agriotes obscurus*, *A. sputator* and *A. lineatus*. The field application rate of 30 kg ha^-1^ for the AgriMet-Granule and 1×10^11^ conidia ha^-1^ for the AgriMet-Dry Product was increased exponentially because no information on the lethal effects was previously available. The Kaplan-Meier survival curves in Figure 14 show, that the mortality caused by the *M. brunneum* formulations was very low, although the field application rates have been increased up to 100 times. Only the AgriMet-Dry Product resulted in a noteworthy reduction of survival probability of *A. obscurus* with significant differences between the application rates (log-rank test, *p* < 0.0001). The field application rate of 1×10^11^ conidia ha^-1^ killed almost no larva of *A. obscurus* and reduced the survival probability only to 98 % after 59 days. A tenfold increase of the application rate to 1×10^12^ conidia ha^-1^ reduced the survival probability to 63 % after 77 days, which differed significantly from the field application rate (Bonferroni, *p* < 0.01). After 66 days of incubation with an application rate of 5×10^12^ conidia ha^-1^ of the AgriMet-Dry Product the survival probability of *A. obscurus* was at 28 % and differed significantly to the lower ones (Bonferroni, 1×10^11^ conidia ha^-1^ = p < 0.0001, 1×10^12^ conidia ha^-1^ = p < 0.01). The LT_50_ of this concentration was 54 days. The highest application rate of 1×10^13^ conidia ha^-1^ led to the lowest survival probability of 18 % after 63 days with a LT_50_ of 45.5 days.

**Figure 7:**
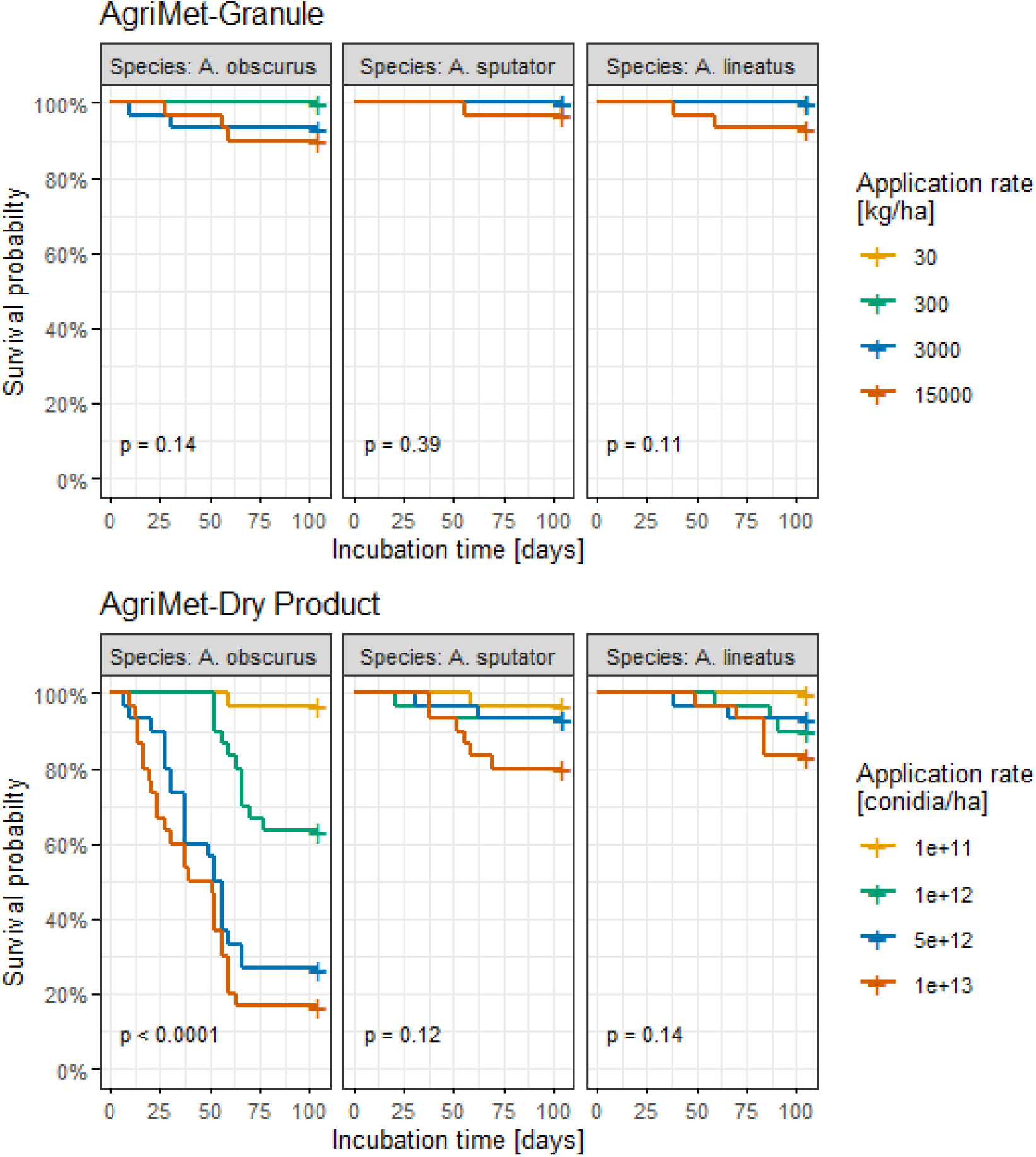
Overall survival probability in percent (Kaplan-Meier-Analysis) of the wireworm species *Agriotes obscurus*, *A. sputator* and *A. lineatus* incubated with four different application rates of the *Metarhizium brunneum* formulations AgriMet-Granule and AgriMet-Dry Product over 105 days under laboratory conditions in small cans filled with soil at 25 °C and darkness. The statistic of the shown *p*-values is based on the log-rank test (*α* = 0.05) and indicates differences between the application rates within a formulation and the respective wireworm species.

However, there was no significant difference between the two highest application rates of the AgriMet-Dry Product. The lethal effect of the AgriMet-Dry Product against *A. sputator* and *A. lineatus* was much weaker and only the highest application rate of 1×10^13^ conidia ha^-1^ was able to reduce the survival probability to a maximum of 80 % after 70 days for *A. sputator* and 82 % after 84 days for *A. lineatus*. The lower application rates resulted in only a few infected larvae of *A. sputator* and *A. lineatus*. The bioassay with the AgriMet-Granule against *A. obscurus*, *A. sputator* and *A. lineatus* resulted in no killed larva at the field application rate of 30 kg ha^-1^. Only very few larvae with a mycosis at the highest application rate of 15000 kg ha^-1^ were examined, but no differences to the lower application rates could be calculated. Because of the very low mortality caused by the *M. brunneum* formulations the calculation of the lethal concentration, at which 50 % of the individuals died, was only possible for the AgriMet-Dry Product and *A. obscurus*. The LC_50_ value after 105 days of incubation was 2.86×10^12^conidia ha^-1^ (FI 95 % 1.68×10^12^-4.96×10^12^ conidia ha^-1^) and is shown in Table 10.

## Discussion

In this study, the effectiveness of the AgriMet-Granule and the AgriMet-Dry Product against wireworms for the use in potato cultivation were tested in field experiments. A prerequisite for a reliable investigation of pesticide effectiveness in the field is the presence of an adequate pest infestation to ensure realistic conditions (Anonymous 2012a). Across all field trials, the proportion of undamaged potatoes in the untreated controls ranged between 0.3-13.1 % indicating a very high degree of wireworm infestation. The reason for this is probably on the cultivation history, as all field sites used for the experiments were uncultivated fallows that were prepared by mechanical tillage shortly before potato planting. Since the previous dense green was suitable for oviposition of click beetles and larval development was not disturbed by any agricultural measures, a large wireworm population was able to establish (Gough & Evans 1942, Seal et al. 1992, Furlan 1996, Furlan et al. 2009). Biocontrol agents often show lower effectiveness in case of a very high pest incidence and are recommended for low or medium infestation, such as the tested reference product ATTRACAP® (BIOCARE GmbH 2020). In the field trials performed in this study, ATTRACAP® with an application rate of 30 kg ha^-1^ reduced the tuber damage only by 2.1-11.3 % compared to the untreated control. The effectiveness of the AgriMet-Granule (30-60 kg ha^-1^) and the AgriMet-Dry Product (1-2×10^11^ conidia ha^-1^) was comparable to ATTRACAP® depending on the respective year, but overall too low for a worthwhile control strategy. Neither the single application of the AgriMet formulations nor the combined application of both could increase the effectiveness above 14 % compared to the untreated control. Previously approved chemical insecticides, such as Fipronil, applied in potatoes achieved an effectiveness of 50-85 %, which would also be desirable for the AgriMet formulations (Kuhar & Alvarez 2008, Ladurner et al. 2009). The low effectiveness is probably due to a complex combination of various factors, which influence the control potential of *M. brunneum* in the soil. The following discussion focuses on these factors and suggestions on how a successful wireworm control strategy based on *M. brunneum* might be implemented.

In general, the AgriMet formulations were intended to ensure pest reduction by inoculative biocontrol. For this, the biocontrol agent *Metarhizium brunneum* should proliferate after application, form conidia in the soil and kill wireworms by initiating the lethal infection cycle (Hajek & St Leger 1994, Roy et al. 2006, Vega et al. 2012). The level of wireworm susceptibility is closely related to the conidia concentration (Quesada-Moraga et al. 2006, Gabarty et al. 2014). The first factor in this context was the appropriate application rate of the AgriMet-Granule and AgriMet-Dry Product, which was investigated under standardised conditions in the greenhouse and laboratory. Using the field application rates in the greenhouse experiment resulted in a slightly better effectiveness of 21.9-25.4 %. In addition to the constant temperature in the pots, regular moistening by watering might have favoured fungal proliferation and thereby enhanced the effectiveness (Dimbi et al. 2004, Ekesi et al. 2003, Arzumanov et al. 2005). An acceptable effectiveness in the greenhouse of up to 44.1-53.9 % was achieved by increasing the application rate of each formulation by a factor of one hundred. However, the reduced tuber damage in the greenhouse was not the result of killed wireworms, since no significant differences in the number of recaptured larvae compared to the untreated control were observed. Therefore, the laboratory experiment was designed to provide information on the lethal potential of the formulations, because reducing wireworms was the overall goal of the control strategy.

The testing of the AgriMet formulations in the laboratory revealed that even an application rate of 15,000 kg ha^-1^ of the AgriMet-Granule resulted only in a 2-10 % decrease in survival probability for all tested wireworm species. This indicated a lack of sufficient conidia formation and related lethal effectiveness. In contrast, the AgriMet-Dry Product with application rates of 5×10^12^-1×10^13^ conidia ha^-1^ reduced the survival probability of *A. obscurus* by 82 %, which underlined its control potential. However, in contrast to the greenhouse and field experiments, larvae in the laboratory bioassay were exposed to the AgriMet-Dry Product in very narrow space without the possibility to avoid fungal infested areas. Based on the inoculative biocontrol approach, extensive proliferation of *M. brunneum* in the potato ridge is required to provide the lethal potential examined in the laboratory experiments. The abundance of *Metarhizium* spp. in the potato ridge during field trials, described as the colony-forming units (CFU) per g soil, reached a maximum of 959.9 CFU g^-1^ by the combined application of the AgriMet-Dry Product (1×10^11^ conidia ha^-1^) and the AgriMet-Granule (30 kg ha^-1^) in 2018. None treatment surpassed 200 CFU g^-1^ soil in 2019 and 2020. Since the desired proliferation of *Metarhizium* spp. in the potato ridge could only be detected to a limited extent, the AgriMet formulations do not seem to be suited for the desired inoculative control strategy and several reasons are worth considering. First, the area of soil sampling may not match with the area in which the formulations were applied. Immenroth (not published) verified the distribution of the AgriMet-Granule evenly and close around the seed potatoes. The simple spray application of the AgriMet-Dry Product with the potato seed dressing during planting also accomplished a surface distribution of infectious *M. brunneum* conidia in the area of the seed potatoes (Matthews 2008). As a result, an error in soil sampling can be excluded for the low *Metarhizium* spp. concentration in the potato ridge. Second, the soil properties may have had an impact on the fungal proliferation after mycoinsecticide application. Since the analysed properties of the light loamy-sandy soils of the different field sites were comparable and did not show any striking values, this aspect can also be excluded. It is, however, more likely that the application rate might be responsible for the low CFU’s g^-1^ soil recovered during the field trials. Consistent with the results presented here, a previous experiment by Rogge et al. (2017), using *M. brunneum* formulated as fungal colonised barley kernels, showed that the application of 1×10^12^ conidia ha^-1^ resulted in 153-456 CFU g^-1^ substrate. With this treatment, most of the tested wireworms survived. However, the authors reported that an application rate of 1×10^15^ conidia ha^-1^ led to 182,945 CFU g^-1^ substrate. This concentration of approximately 1×10^6^ CFU g^-1^ soil significantly reduced the number of tested wireworms and should therefore be used as benchmark for inoculative biocontrol strategies. Thus, the application rate may have had an influence on the low effectiveness of the AgriMet formulations during the field trials due to the insufficient proliferation of *M. brunneum* and related lack of infectious conidia. Despite the promising results of the AgriMet-Granule and AgriMet-Dry Product at very high application rates in the greenhouse, 3000 kg ha^-1^ or rather 1×10^13^ conidia ha^-1^ are not feasible for particle use, since the application technology has limitations (personal communication, Lehner Maschinenbau GmbH, 2020). Moreover, the cost for the formulations would exceed the additional revenue from more marketable potatoes. However, the evaluation of the application rate has to be considered with caution, as no consistent product stability was given.

Product stability is an important formulation property that determines, among other factors, storability. *M. brunneum* needs to be remain viable during storage without loss of activity and degradation of the desired formulations properties (Jones & Burges 1998). However, both the AgriMet-Granule and the AgriMet-Dry Product show deficits in terms of product stability during storage. For the AgriMet-Dry Product, a concentration of 1.25×10^9^ CFU g^-1^ was reported by the manufactures and 5.6-9.6×10^8^ CFU g^-1^ was finally assessed. The small loss of vitality could be due to the heterogeneous particle size of the powder and the small quantity used for the quality control. The AgriMet-Granule also showed clear deficits in storage stability four weeks after shipment. A vitality loss of up to 20 % in individual granule batches was assessed, compared to the manufacture specifications, indicating a critical factor influencing the effectiveness. Based on the results of the quality control, biological activity was expected on average only at 75 % of the respective quantities applied, which might be another explanation for the low CFU values in the field trials. Since the outgrowth rate of the reference product ATTRACAP® ranged between 97-100 %, encapsulation of the fungal biocontrol agent seems to provide better protection during storage compared to the outer coating of the AgriMet-Granule. Additional coatings on the AgriMet-Granule could be helpful to protect the fungus from physical or chemical influences and thus improve product stability (Jones & Burges 1998). Besides the preserved vitality of the organism during storage and shipment, the fungus must be able to proliferate under the environmental conditions at the target site. Therefore, abiotic factors must also be considered when evaluating the effectiveness to account for the ecological requirements of *M. brunneum*.

Granule formulations of entomopathogenic fungi with cereal grains are advantageous in principle because they provide a suitable substrate for fungal growth and prolong persistence in the soil (Burges 1998). Since only 5.89×10^4^ conidia grain^-1^ were determined on the surface of the AgriMet-Granule, the thin *M. brunneum* layer on the surface of the autoclaved millet must first proliferate for the formation of sufficient fungal material after application. Growth, germination and related virulence of *Metarhizium* spp. are temperature-dependent with an optimum of 25-30 °C, whereas lower (15 °C) or higher (35 °C) temperatures influence the development negatively (Dimbi et al. 2004, Ekesi et al. 1999, Bugeme et al. 2009, Tumuhaise et al. 2018). Bernhardt (not published) verified the mentioned temperature conditions for the *M. brunneum* isolate JKI-BI-1450 used for the AgriMet formulations. At the beginning of all field trials, the soil temperature in the potato ridge showed a daily mean of 9.8-14.7 °C. Soil warmed up very slowly, reaching an average temperature of 17.1 °C. A daily mean temperature of 25 °C was determined on a single day in the end of July 2019. Overall, the soil temperature in the potato ridge was too low for optimal growth and germination of isolate JKI-BI-1450, especially during the application in April/May. This was confirmed by the quality control of the AgriMet-Granule in field soil, which resulted in no visible fungal growth or sporulation after incubation at 15 °C for 14 days. Crucial problems arise from the very slow growth of the *M. brunneum* isolate JKI-BI-1450 on the AgriMet-Granule at low soil temperatures. First, seasonal activity of wireworms requires lethal potential by *M. brunneum* shortly after application. The wireworms move into upper soil layers in early spring and autumn preferring foraging at suitable soil moisture and temperature (Langenbuch 1932, Brian et al. 1947, Burrage et al. 1963, Parker & Howard 2001). Summer drought as well as moulting can decrease wireworm activity in the upper soil layer in between these periods, reducing the likelihood of contact with formulated *M. brunneum* (Evans & Gough 1942, Furlan 1998, Staley et al. 2007). Second, larval death can take 41-80 days after contact with fungal conidia, depending on the respective isolate and wireworm species (Brandl et al. 2017). Therefore, infection by *M. brunneum* later in the potato growing season would not prevent damage to daughter tubers (Hack et al. 1993). Third, until *M. brunneum* proliferation on the AgriMet-Granule occurs, the autoclaved millet grain serves as nutrient-rich substrate for ubiquitous soil-borne fungi using accessible carbon sources for metabolism and growth (Geervani & Eggum 1989, Dighton 2007, Kennedy 1994, Deacon 2013). Especially opportunistic saprophytic fungi from agricultural soils are adapted to low temperatures and might compete for the autoclaved millet after application (Hajek 1997, Pietikäinen et al. 2005, Bárcena-Morena et al. 2009). The decomposition of the coated autoclaved millet grain might prevent the development of the isolate JKI-BI-1450. Thus, wireworm control with the AgriMet-Granule cannot be successful at the recorded temperatures. However, even the incubation of the AgriMet-Granule at 25 °C for 14 days only led to a minimal increase in conidia concentration to 7.1×10^5^ conidia grain^-1^. This might explain the lack of lethal potential in the greenhouse and laboratory tests, since insufficient amounts of conidia are formed in soil to kill wireworms. However, the AgriMet-Dry Product is also negatively affected by the low temperatures during application. The spray application should directly hit the wireworms through droplets or colonise organic material, but fungal germination should be unlikely under the conditions recorded. Thus, temperature in the potato ridge is another critical factor influencing the effectiveness of the AgriMet formulations in the field. Effectiveness of mycoinsecticide formulations against wireworms may be improved by using isolates of the genus *Metarhizium* from other geographic regions that show greater thermotolerance for low temperatures. For example, Isolate ARSEF 4343 originating from Macquarie Island close to the Antarctica was cold-active even at 5 °C and might be better suited for the use in potato cultivation (Fernandes et al. 2008). However, the thermotolerance of an isolate must be in proportion to its virulence, which leads to the next point of discussion.

Usually, several wireworm species occur simultaneously in one field because larval development takes 3-5 years and the populations overlap (Subklew 1935, Miles 1942, Klausnitzer 1994, Furlan 1998). Thus, a universal and effective application of the AgriMet formulations in potato cultivation throughout Germany requires a high virulence of formulated *M. brunneum* isolate JKI-BI-1450 against the most important wireworm species. However, *A. obscurus*, *A. sputator* and *A. lineatus*, exposed to the AgriMet-Dry Product in the laboratory bioassay, showed significant differences in mortality. While the survival probability of *A. obscurus* was reduced by 80 %, the survival probability of *A. sputator* and *A. lineatus* decreased by only 20 %, indicating selectivity of isolate JKI-BI-1450. Bernhardt et al. (2019) confirmed the selectivity of isolate JKI-BI-1450 against *A. obscurus* in a standardised laboratory dip test. As *A. lineatus* was the most abundant wireworm species during the performed field trials and isolate JKI-BI-1450 showed only low virulence against it, no relevant control potential could be expected by the application of the AgriMet formulations in the field. The specific virulence of *M. brunneum* isolates intended for wireworm control poses a crucial problem in practical use. Lehmhus (2019) reported a diverse wireworm species composition on German potato fields varying by region. The most common species are *A. obscurus*, *A. sputator* and *A. lineatus*, but further species of agronomic importance like *A. ustulatus* and *A. sordidus* occurred in the south/southwest of Germany. Lehmhus (2019) postulated that the change to a warmer climate might advance the distribution of new, economic important species. It is therefore unlikely that a mycoinsecticide containing a single biocontrol agent is capable to control the diverse pest complex of wireworms.

Several factors such as the application rate, soil temperature and isolate selectivity might explain the low or missing effectiveness of the AgriMet formulations in the field. Consequently, the recorded reduction of tuber damage during the field and greenhouse experiments cannot be attributed to the reduction of wireworms alone. The effect of the AgriMet formulations in the field and greenhouse might probably be the result of behavioural changes of wireworms after fungal infection. Previous studies with the desert locust *Schistocerca gregaria* and the grasshopper *Zonocerus variegatus* demonstrated a reduction of feeding after infection with *M. flavoviride* (Moore et al. 1992, Thomas et al. 1997). The pollen consumption of the flower thrips *Megalurothrips sjostedti* was also reduced by an infection with *M. anisopliae* (Ekesi et al. 1999). A sublethal infection of wireworms may have influenced the foraging behaviour and thereby resulted in a reduction of tuber damage. Another partial reason for the field trial results could be the spatial distribution of wireworms at the field site and the associated experimental design. Because click beetles can lay eggs in multiple clusters during oviposition, the occurrence of larvae in the field is not uniform (Furlan 1996, Sufyan et al. 2014). Wireworm sampling in each plot confirmed a patchy distribution, and minor tuber damage, regardless of treatment, was observed primarily in plots where very few or no wireworms were found. These individual plots may have influenced the significance of the statistical comparison of the mean proportion of undamaged potatoes. Reliable data could be generated by using more blocks/repetitions in the experimental design, reducing the weighting of each plot. Another option could be the systematic placement of blocks at the field site, taking into account the distribution of wireworms. The use of excluded or adjacent untreated controls could be useful to evaluate the effectiveness of treatments without bias from wireworm distribution (Anonymous 2012a).

In conclusion, none of the AgriMet formulations based on the biocontrol agent *M. brunneum* improved tuber protection compared to ATTRACAP® after field application. Insufficient product stability, low soil temperatures during spring application and the limited virulence of the formulated isolate JKI-BI-1450 against *A. lineatus* and *A. sputator* may have had a decisive influence on the control potential of wireworms. The AgriMet-Granule and the AgriMet-Dry Product need be optimised based on these influencing factors to ensure a meaningful field use. Future field research on the effectiveness should be based on knowledge of the detailed composition and spatial distribution of wireworm species to avoid bias due to the experimental design. Only reliable effectiveness data of field trials can convince agricultural practitioners to use *M. brunneum* against wireworms in potato cultivation and promote sustainable agriculture through biocontrol.

